# Colonic inflammation modulates the intestinal circadian landscape

**DOI:** 10.1101/2025.02.20.639289

**Authors:** Thomas D Butler, Polly Downton, Suzanna H Dickson, Andrea Luengas-Martinez, Devin A Simpkins, Isabel Khoo, Sarah Veal, Alexander C West, Antony D Adamson, David A Bechtold, John T McLaughlin, Julie E Gibbs

## Abstract

Inflammatory bowel disease (IBD) is a chronic inflammatory condition that carries significant morbidity. The circadian (24h) clock regulates many aspects of immunity and circadian disruption is implicated in inflammatory disease. We examined the impact of acute dextran sulphate sodium-induced colitis on the intestinal circadian landscape and investigated consequences of intestinal epithelial cell (IEC) clock disruption on immune function in health and under inflammatory stress. Within the inflamed colon, we observed attenuated IEC clock function. Furthermore, *de novo* oscillations in numbers of colonic regulatory T cells emerged, associated with diurnal variation in proliferation and activation markers. IEC-specific deletion of core clock gene *Bmal1* drove alterations in the rhythmic colonic transcriptome (impacting key immune pathways) and marked damping of microbiota rhythmicity. IEC-specific *Bmal1* deletion did not impact colitis severity. These results highlight the impact of colonic inflammation on circadian processes and suggest that circadian logic could be applied in IBD treatment.

## Introduction

Inflammatory bowel disease (IBD) is characterised by chronic intestinal inflammation interspersed with flares of acute inflammation, leading to significant morbidity. Gut inflammation is driven by disruption of intestinal epithelial cell (IEC) function, breakdown of host-microbiome interaction and subsequent expansion of inflammatory effectors into the lamina propria layer ^1^. Despite significant advances in immunosuppressive medication, life-changing surgery is frequently required, necessitating further work to understand drivers of IBD.

Intrinsic molecular clocks, found in nearly all lifeforms, have evolved to drive 24h circadian rhythms in physiology and behaviour aligned to the Earth’s rotation. The mammalian circadian clock is driven by molecular feedback loops involving transcriptional activators BMAL1/CLOCK and transcriptional repressors PERIOD, CRYPTOCHROME and NR1D1 (REV-ERBα). The circadian clock is vital for effective immunity and regulates a plethora of immune functions including leukocyte trafficking, cytokine production and response to pathogens ^2^. Within the gut, IECs contain a functional clock that responds to rhythmic signals including hormones, cytokines and neuropeptides ^3–5^. Additionally, the gut clock is responsive to extrinsic signals derived from the diet and gut microbiota, both of which exhibit 24h rhythmic patterning. This results in temporal co-ordination of multiple intestinal processes including antigen presentation, antimicrobial peptide production, host-microbiome interaction and nutrient absorption ^6–9^. The IEC therefore sits at the interface between microbiome and host immunity and its robust clock is sensitive to multiple environmental signals. Data is beginning to emerge highlighting the importance of local tissue clocks in regulating gut inflammation, ^10,11^ but little is known about the impact of gut inflammation on IEC rhythmicity.

Circadian disruption is implicated in pathogenesis of chronic inflammatory conditions, such as rheumatoid arthritis and asthma ^12^. In people with IBD, inflamed colonic tissue has damped core clock gene expression, and there are growing genetic associations of clock components and IBD phenotype ^13,14^. Furthermore, in mice, environmental and genetic circadian disruption has been shown to modulate colitis severity ^15–18^.

Here we investigate the consequences of inflammation on the intestinal circadian landscape in mice and study the impact of IEC clock disruption on immune function in health and colitis. We demonstrate that mice with DSS-induced colitis exhibit marked damping of the IEC clock early in disease. Through the generation of transgenic mice with cell-specific deletion of core clock gene *Bmal1*, we mapped consequences of IEC clock disruption on 24h rhythms in the colonic transcriptome and microbiota, revealing significant temporal re-organisation. Surprisingly, these mice did not demonstrate altered susceptibility to acute DSS colitis, in contrast to recently published work ^10,11^. Despite damping of the IEC clock, mice with colitis have striking *de novo* rhythmicity in lamina propria leukocyte numbers including regulatory T cells (Tregs), which also exhibit diurnal variation in activity within the inflamed gut. Together, these data highlight the importance of functional circadian clocks for intestinal health, and reveal the impact of colonic inflammation on circadian processes.

## Results

### Acute DSS-induced colitis exhibits circadian characteristics

To effectively examine the impact of colitis on the intestinal circadian landscape, we developed a robust experimental approach whereby timed sample collection occurs after a defined and consistent period (7 days) of DSS exposure. C57BL/6 mice were treated with 2.5% DSS for 7 days, initiated at zeitgeber time (ZT)0, ZT6, ZT12 or ZT18 (where ZT0 represents lights on and ZT12 represents lights off). On day 7, tissues were harvested at ZT0, ZT6, ZT12 and ZT18 (**Figure 1A**). DSS treatment induced colitis in each experimental group, and disease severity was not impacted by time of initiation, as observed by weight loss, colon length, severity score and histology score (**Figure 1B-F**). In contrast, when DSS exposure was initiated at a single time point and tissue sampled sequentially at 6-hourly intervals from day 7 (ZT0) to day 8 (ZT0), progressive weight loss and progressive non-rhythmic inflammatory activity was observed (**Figure S1A-S1B**), indicating that staggered DSS initiation and tissue collection is a robust methodology to standardise DSS exposure time for circadian studies.

**Figure 1.**
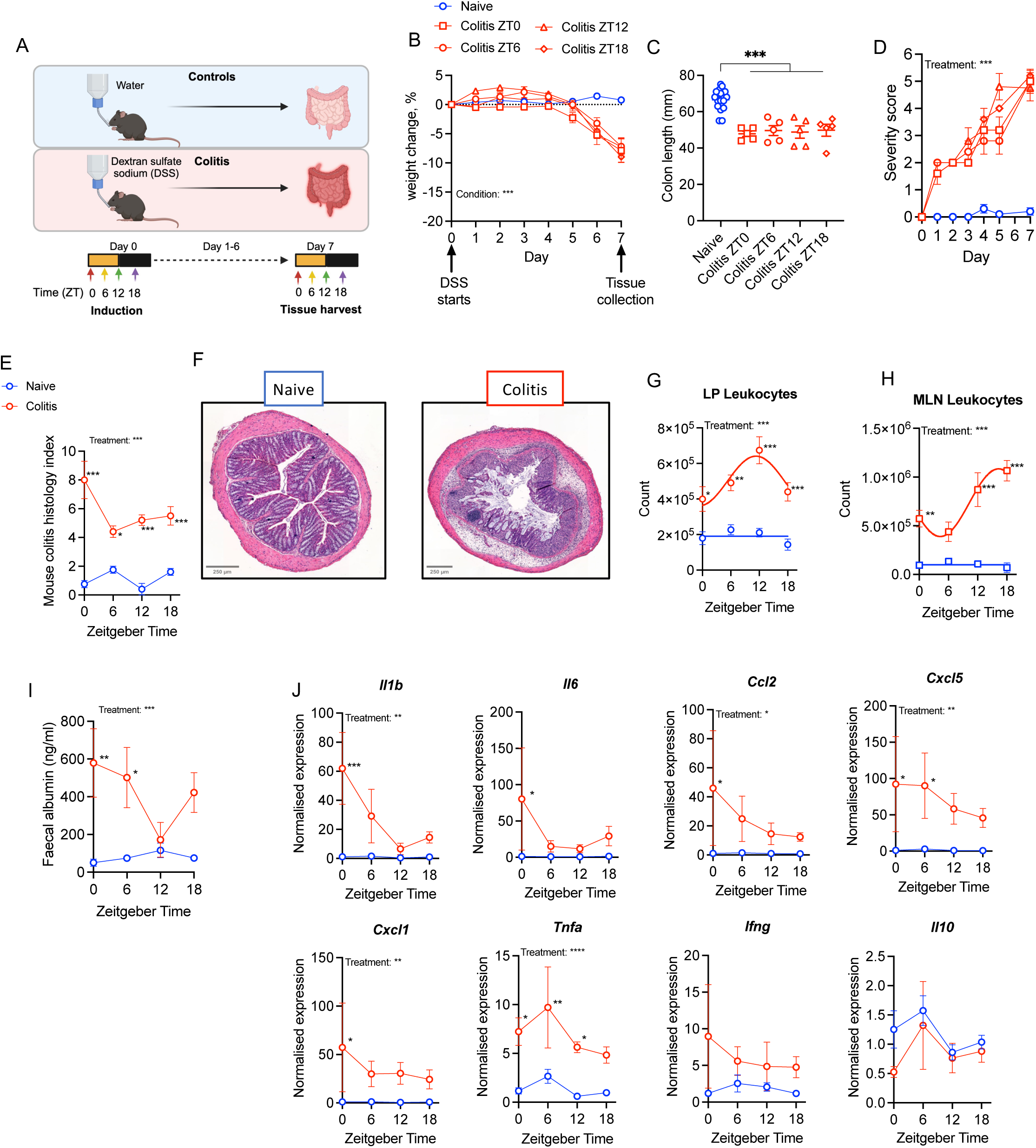
Acute DSS induced colitis exhibits circadian characteristics: **(A)** A schematic of the experimental design for acute dextran sulphate sodium (DSS) induced colitis. **(B)** Percentage weight change of mice exposed to water or 2.5% DSS initiated at Zeitgeber Time (ZT)0, ZT6, ZT12 or ZT18, where ZT0 represents lights on, and ZT12 represents lights off. Data were normalised to day 0 weight (DSS treated, n=5/timepoint; Naïve controls, n=5/timepoint (data pooled into one group). **(C)** Colon length measured from distal colon to anus, n=5/group, controls grouped. **(D)** Daily severity score, n=5/group, controls grouped. **(E)** Mouse colitis histology index scored on mid-colon at day 7, n=5/timepoint/treatment. **(F)** Representative images of H+E-stained colon. **(G)** Number of lamina propria (LP) leukocytes (live CD45^+^), assessed by flow cytometry, n=5/timepoint/treatment. **(H)** Number of mesenteric lymph node (MLN) leukocytes (live CD45^+^), assessed by flow cytometry n=5/timepoint/treatment. **(I)** Intestinal barrier function measured by faecal albumin ELISA at day 7 of DSS treatment, n=4-5/timepoint/treatment. **(J)** qPCR data from colon samples collected at day 7, represented as fold change in ΔΔCt values, using naïve ZT0 as referent population and *B-actin* as housekeeping gene. N=4-5/timepoint/treatment. Statistics: (B,D and J) Two-way ANOVA with multiple comparisons (Tukey). (C) One-way ANOVA with multiple comparisons (Dunnett). (E and I) Two-way ANOVA with multiple comparisons (Šídák). (G and H) Two-way ANOVA with multiple comparisons (Šídák) and nonlinear regression to compare whether best fit is given by horizontal line or sine wave with nonzero baseline, constraints: wavelength = 24 hours; amplitude > 0.

Colitis in mice and humans is associated with localised pro-inflammatory response, infiltration of multiple leukocyte subsets (including T cell subsets) to the lamina propria (LP) and increased intestinal permeability. We examined whether these disease manifestations exhibited time of day variation. Flow cytometric analysis of leukocyte numbers (CD45^+^) within the colonic LP and gut draining mesenteric lymph nodes (MLN) across the 24h day demonstrated *de novo* rhythmicity in DSS-treated mice, with LP leukocytes peaking at ZT12 and MLN leukocytes peaking at ZT18 (**Figure 1G and H**). The number of CD45^neg^ LP cells (likely endothelial cells and stromal cells) were not rhythmic in either condition (**Figure S1C**), demonstrating that this emergent rhythmicity in CD45^+^ LP cells in DSS-treated mice is independent of tissue processing.

IEC barrier function (assessed by fecal albumin content) was significantly impaired after DSS exposure, and permeability varied by time of day (**Figure 1I**). In DSS-treated mice, fecal albumin was significantly elevated at ZT0 and ZT6 (animal’s rest phase) but not significantly different from water controls at ZT12 and ZT18 (animal’s active phase). This is unlikely to be due to a change in feeding behaviour; whilst total daily food intake decreased over the course of DSS treatment, temporal calorie intake was similar between mice treated with DSS and controls, with the majority of feeding expectedly occurring during the night (**Figure S1D-S1E**).

Colonic tissue from mice with DSS colitis demonstrated significantly higher expression of pro-inflammatory cytokines *Il1b, Il6, Ccl2* and *Cxcl1* at ZT0, but no significant difference at ZT6, ZT12 or ZT18 compared to water controls (**Figure 1J**). In mice with DSS colitis, colonic *Cxcl5* expression was significantly higher during the light phase at ZT0 and ZT6, but not ZT12 and ZT18, and *Tnfa* was significantly elevated at all timepoints except ZT18 (**Figure 1J**). Colonic *Ifng* and *Il10* expression was not affected by DSS treatment. Together, this shows the magnitude of colonic inflammation varied across the day and the intestinal environment may be more pro-inflammatory as mice enter their rest phase.

### Alteration of the IEC clock in response to localised inflammation

In DSS-treated mice, *de novo* rhythms in IEC barrier function and leukocyte number suggest an important role for the IEC clock in gut inflammation. Conversely, tissue inflammation is associated with repression of core components of the molecular clock, including REV-ERBα ^19,20^. Core clock gene expression in IECs was significantly damped after 7 days of DSS treatment, and although expression of *Bmal1*, *Reverba*, *Per2* and *Cry1* remained statistically rhythmic, amplitude was markedly attenuated (**Figure 2A**). To understand the temporal association between DSS initiation and clock disruption, mice were generated which express a luciferase-tagged *Reverbα* specifically within IECs (IEC-*Reverbα*^luc^). As expected, mice expressing the IEC-specific luciferase tagged gene demonstrated detectable *in vivo* bioluminescence localised within the intestinal region at ZT8 (**Figure 2B**). Quantification of this signal at ZT8 (peak of colonic *Reverbα* expression ^21^) and ZT0 confirmed that in healthy animals IEC *Reverbα* expression exhibited robust diurnal rhythmicity (**Figure 2C**). After only 1 day of DSS treatment, diurnal variability in *Reverbα* was significantly attenuated, with further attenuation on day 3 (**Figure 2D**). Thus, the IEC clock is impaired early in DSS colitis, and this may contribute to the resulting inflammatory cascade and phenotypic colitis induction.

**Figure 2.**
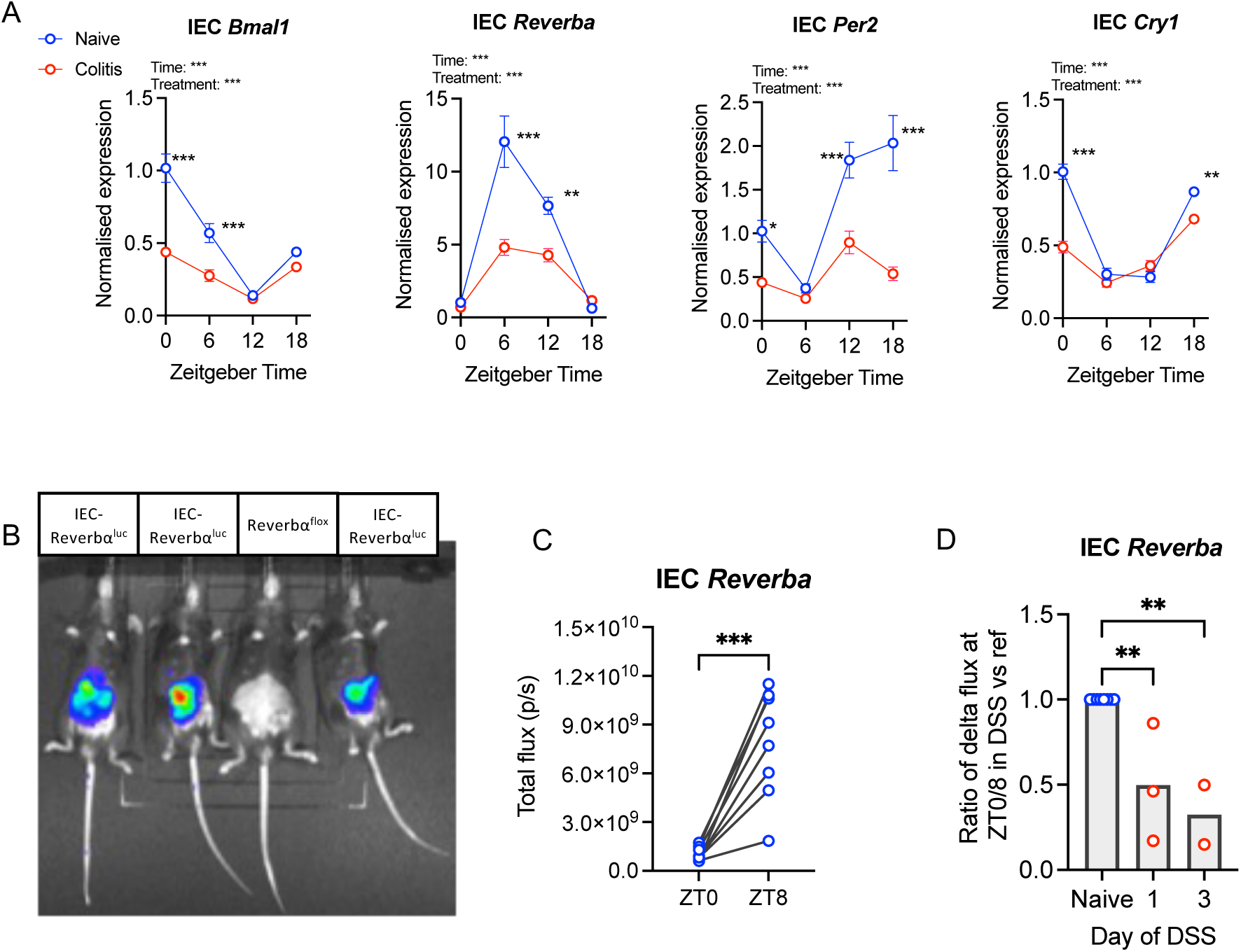
Alteration of the IEC clock in response to localised inflammation: **(A)** Colonic intestinal epithelial cell (IEC) clock gene expression after 7 days of dextran sulphate sodium (DSS) or water, represented as fold change in ΔΔCt values, using naïve zeitgeber time (ZT)0 group as referent population and *Gapdh* as housekeeping gene. n=5/timepoint/treatment. **(B)** Representative *in vivo* imaging system (IVIS) image demonstrating bioluminescent signal from IEC-Reverbα^luc^ reporter mice but not Reverbα^flox^ controls. **(C)** Bioluminescent flux recorded from naïve IEC-Reverbα^luc^ reporter mice each imaged at ZT0 and ZT8, n=8. **(D)** A ratio of the fold change of bioluminescent flux on day 1 (n=3) and day 3 (n=2) of DSS versus referent naïve ZT0:ZT8 flux. Statistics: (A) Two-way ANOVA with multiple comparisons (Tukey). (C) Paired two-tailed t test. (D) One-way ANOVA with multiple comparisons (Dunnett).

### IEC-specific *Bmal1* deletion alters the colonic transcriptome

To address the role of the IEC clock in regulation of immunity and inflammation, IEC-*Bmal1*^−/−^ mice were generated, which lack the core clock gene *Bmal1* in Villin-expressing cells (**Figure 3A and B, and Figure S2A**). BMAL1 was targeted due to its non-redundant core clock gene function ^22^ and more severe colitis described in global *Bmal1*^−/−^ mice ^16^. Villin is highly expressed in IECs and is commonly used to specifically target this population for gene deletion ^23,24^. Using behavioural studies and EchoMRI we demonstrated that naïve IEC-*Bmal1*^−/−^ mice and *Bmal1*^flox^ controls showed no difference in circadian period (**Figure 3C and Figure S2B-S2C**); timing and volume of calorie intake (**Figure 3D**); or body composition (**Figure 3E**). Furthermore, IEC-*Bmal1*^−/−^ mice treated with 16 weeks of 60% high-fat diet were significantly more resistant to diet-induced obesity (reduced weight gain) compared with *Bmal1*^flox^ mice, in keeping with recently published data ^9^ (**Figure S2D-S2E**), supporting successful induction of IEC-specific *Bmal1* deletion. To characterise impact of IEC-specific *Bmal1* deletion on the colonic transcriptome, RNASeq was performed on tissue harvested 6-hourly across the 24h cycle. Of 21,113 transcripts, global analysis (with no consideration of time of sample collection) revealed significant upregulation of 147 (0.7%) and downregulation of 111 (0.5%) genes in IEC-*Bmal1*^−/−^ mice (adjusted p value < 0.05) (**Figure 3F)**. As expected, clock gene expression was significantly impacted, with up-regulation of *Cry1* and *Npas2* and down-regulation of *Nr1d1/2* (encoding *Reverbα*/*β*). Of note, *Cldn8* was significantly upregulated and lost rhythmicity in IEC-*Bmal1*^−/−^ mice (**Figure 3G**). *Cldn8* encodes CLAUDIN-8, a key component of IEC tight junctions implicated in IBD pathogenesis ^25,26^, which prompted us to test barrier permeability (**Figure 3H**). Results demonstrated no significant genotype effect, however naïve IEC-*Bmal1*^−/−^ mice trended towards more pronounced diurnal variation in fecal albumin concentration. Temporal expression of additional genes associated with tight junctions (KEGG 2019 pathway: Tight junction; mmu04530) were also impacted in IEC-*Bmal1*^−/−^ mice (**Figure S2F**).

**Figure 3.**
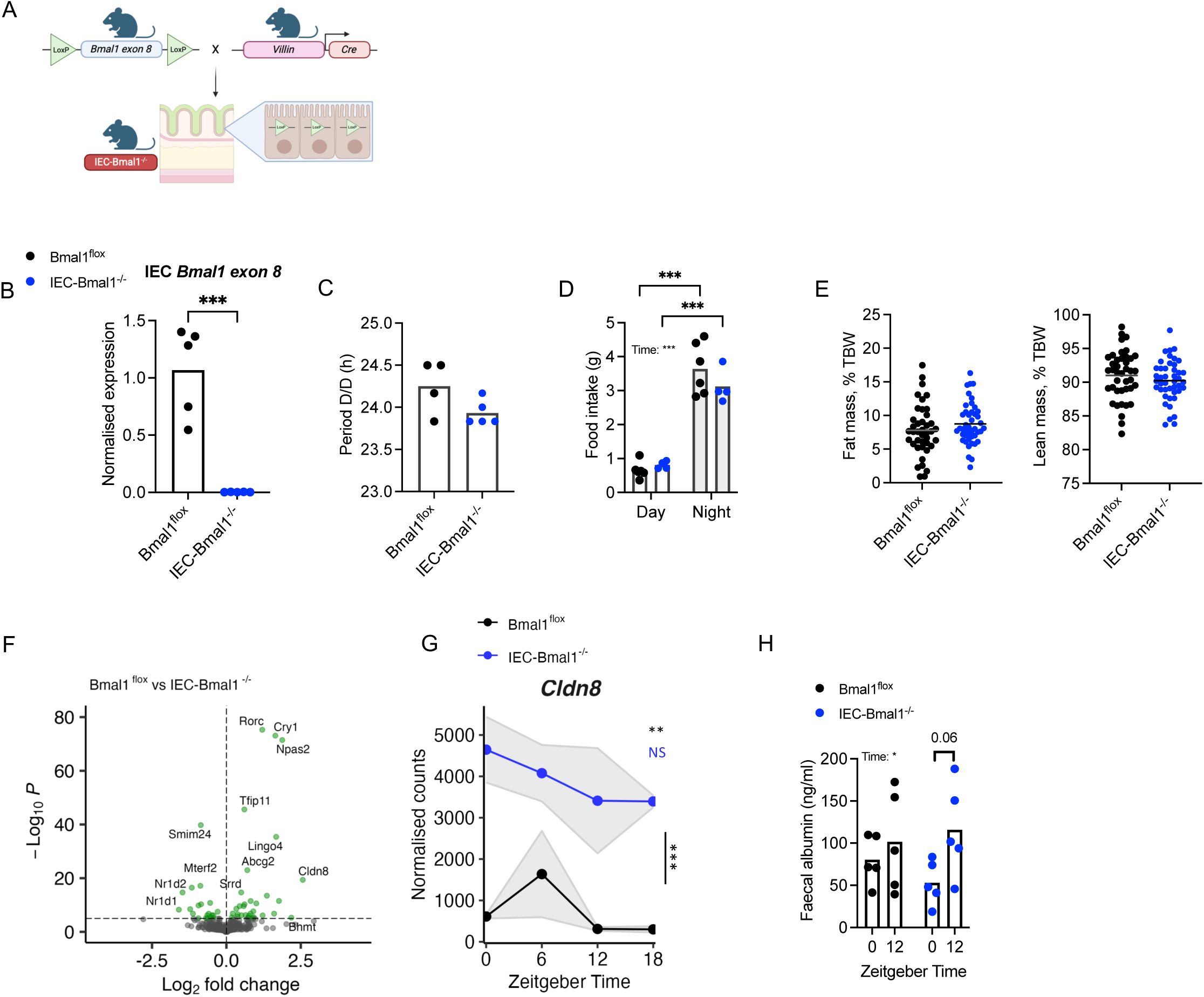
IEC-specific *Bmal1* deletion alters the colonic transcriptome. **(A)** Schematic showing the breeding used to generate *Bmal1*^fl/fl^ *Villin*^Cre/+^ (intestinal epithelial cell (IEC)-*Bmal1*^−/−^) mice. **(B)** Colonic IEC clock gene expression in naïve mice, represented as fold change in ΔΔCt values, using naïve zeitgeber (ZT)0 group as referent population and *Gapdh* as housekeeping gene, n=5/genotype. **(C)** Wheel running periods determined during constant dark conditions, n=4-5/genotype. **(D)** Temporal food intake during the day (ZT0-ZT12) and night (ZT12-ZT0) in single-housed mice, n=4-6/genotype. **(E)** Body composition analysis determined by EchoMRI using mice maintained on normal chow, n=41-45/genotype. **(F)** Volcano plot illustrating differentially expressed (DE) colonic transcripts between naive IEC-*Bmal1*^−/−^ (n=16) and *Bmal1*^flox^ (n=15) mice, Log_2_ fold change cutoff; p-value cutoff, 10e^-5^. **(G)** Temporal expression of *Cldn8* transcripts from RNASeq data. Error bars represent standard error of the mean. **(H)** Intestinal barrier function measured by faecal albumin ELISA after 7 days of DSS or water, n=5/genotype. Statistics: (B, C, E) two-tailed t test; (D, H) Two-way ANOVA with multiple comparisons (Šídák); (G) Two-way ANOVA and JTK_CYCLE.

### IEC-specific *Bmal1* deletion alters the colonic rhythmic transcriptome, rhythmic microbiome and key immune pathways

Transcriptome rhythmicity was assessed using compareRhythms, a robust analysis tool that uses probability measures between genotype groups to assign ‘same’ or differential (‘gained’, ‘lost’, ‘changed’) rhythmicity ^20,27^. Of the 2,995 rhythmic genes identified in the colon, 43% were differentially rhythmic between genotypes, with 1,047 showing loss of rhythmicity in IEC-Bmal1^−/−^ mice, 209 gain of rhythmicity and just 37 assigned a change (**Figure 4A**). As expected, temporal expression of core circadian clock genes was significantly altered in the IEC-*Bmal1*^−/−^ colon, with attenuated amplitude of rhythmicity in *Nr1d1*, *Βmal1*, *Dbp* and *Nfil3* (**Figure 4B**). Thus, whilst IEC-specific *Bmal1* deletion had limited effect on differentially expressed (DE) genes, there is considerable disruption to the rhythmic transcriptome.

**Figure 4.**
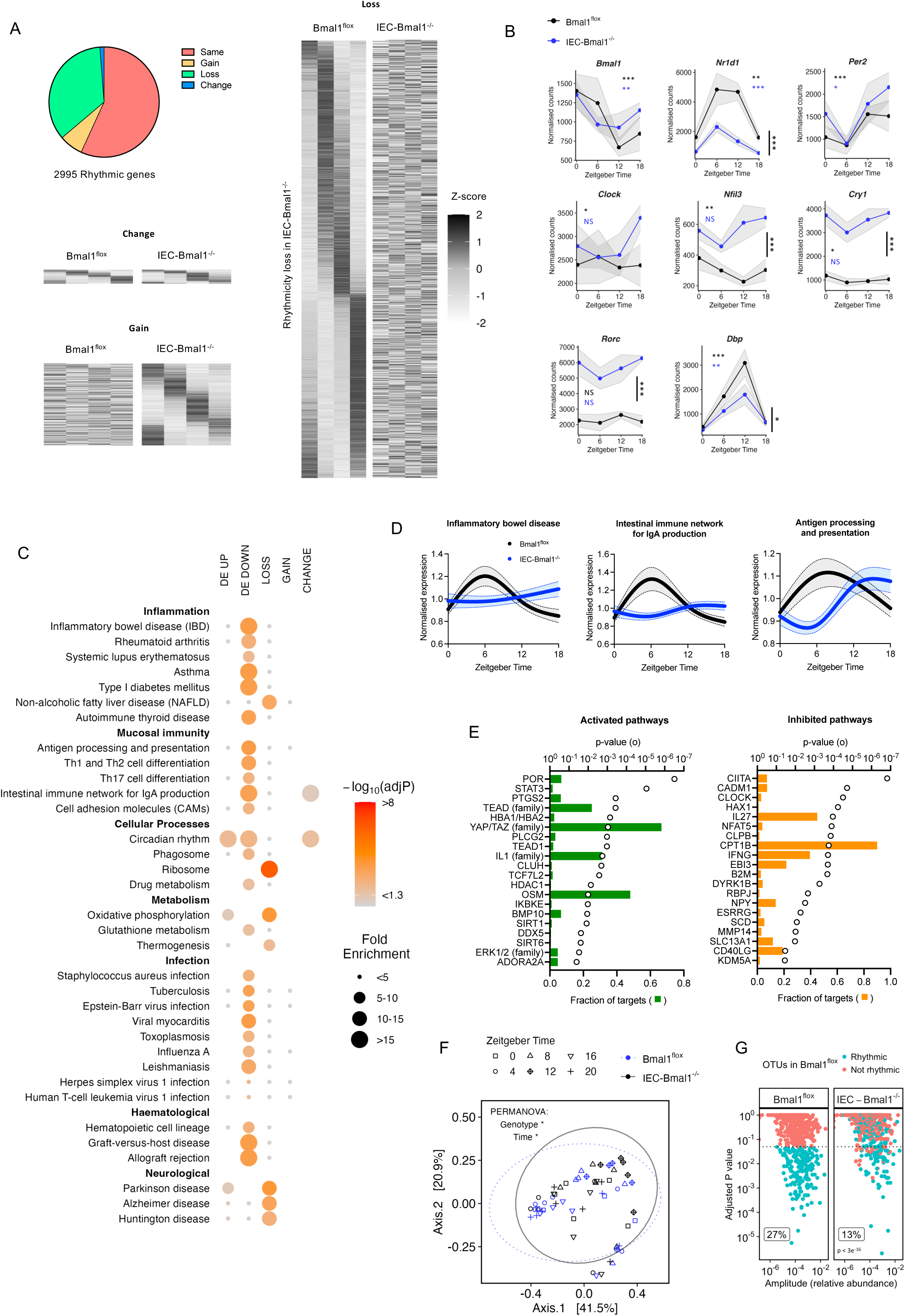
IEC-specific *Bmal1* deletion alters the colonic rhythmic transcriptome, rhythmic microbiome and key immune pathways. **(A)** Differential rhythmicity analysis with compareRhythms categorises rhythmicity in colonic transcript expression in naive IEC-*Bmal1*^−/−^ compared to *Bmal1*^flox^ mice. n=3-4/genotype/timepoint (zeitgeber time (ZT)0, ZT6, ZT12, ZT18). **(B)** Core clock gene transcript expression across time. N=3-4/genotype/timepoint. Error bars represent standard error of the mean. **(C)** Functional pathways enriched (using Enrichr tool and KEGG 2019 mouse database) in gene transcripts with significantly differential expression (DE up/down) or significantly differential rhythmicity (loss/gain/change). Size of spot represents category of fold enrichment, colour represents significance of enrichment. **(D)** Spline plots showing mean normalised expression of all genes from the dataset within a selected pathway (inflammatory bowel disease, mmu05321, 54/62; intestinal immune network for IgA production, mmu04672, 37/43; and antigen processing and presentation, mmu04530, 71/87), error bars represent 95% confidence intervals. **(E)** The top 20 activated and inhibited upstream regulator pathways. Open circle represents significance of enrichment by Ingenuity Pathway Analysis, filled bars represent fraction of downstream targets present in dataset. **(F)** Beta diversity (Bray-Curtis) within 16S microbiome sequencing (n=30 per genotype). **(G)** Rhythmicity in relative abundance of OTUs, n=5/timepoint/genotype. Abbreviations: ZT = zeitgeber time; DE = differential expression; OTU = operational taxonomic unit. Statistics: (A) compareRhythms rhythmicity analysis. (B) Vertical lines represent between genotype significance by two-way ANOVA. Coloured, horizontal asterisks indicate rhythmicity analysis by JTK_CYCLE (NS = not significant). (F) PERMANOVA test. (G) Rhythmicity assessment by JTK_CYCLE; Chi squared test for comparison of rhythmic fraction between genotypes.

Pathway enrichment analysis using Enrichr ^28–30^ was performed on differentially rhythmic genes (loss, gain and change) as well as differentially expressed (DE) genes between genotypes, regardless of time of day (DE up and DE down) (**Figure 4C**). Genes that lost rhythmicity were significantly enriched for ribosomal pathway and oxidative phosphorylation. Transcripts with a change in rhythmicity were enriched for circadian pathways and intestinal immune network for IgA production. Interestingly, downregulated colonic transcripts in IEC-*Bmal1*^−/−^ mice were enriched for pathways related to inflammation, mucosal immunity and infection, such as inflammatory bowel disease (mmu05321), antigen processing and presentation (mmu04612), and intestinal immune network for IgA production (mmu04672) (**Figure 4D**). Genes included in each pathway are listed in **Table S1**. Defects in these pathways have been linked to colitis severity and aberrant host-microbe interactions ^31,32^. Similar pathways were identified as perturbed using Ingenuity Pathway Analysis (IPA), including activation of IL-10 signalling pathways and inhibition of key immune pathways such as major histocompatibility complex (MHC) class II antigen presentation (**Figure S3A-S3B)**. We next identified potential upstream regulators using IPA (**Figure 4E**). Recognised inflammatory mediators such as STAT3 and the IL1 family were activated in IEC-*Bmal1*^−/−^ mice, and inhibited pathways included IFNγ, involved in multiple inflammatory signalling pathways including JAK-STAT, and CIITA, a master regulator of major histocompatibility complex (MHC) expression, associated with microbiome regulation ^7,33^. These data highlight the influential role of *Bmal1* within IECs across multiple functions and pathways, including modulation of immunity, inflammation and the microbiome, and suggest disruption of *Bmal1* function influences intestinal homeostasis and may modulate propagation of intestinal inflammation.

Given the important role of IEC function in host-microbiome crosstalk ^34^ and our observed disruption of pathways involving microbiome sampling and processing in IEC-*Bmal1*^−/−^ mice, we monitored the effect of IEC-specific *Bmal1* deletion on circadian rhythms in gut bacterial composition using 16S V4 rRNA sequencing of fecal pellets. In both genotypes, alpha diversity was 24h rhythmic, peaking at ZT16 during the dark active phase (**Figure S3C**). Beta diversity significantly varied by time and also showed a small but significant difference between genotypes (**Figure 4F**). Across all samples, 717 operational taxonomic units (OTU) were identified, spanning 10 phyla. In naïve *Bmal1*^flox^ controls, 27% (191/717) of OTUs demonstrated significant daily rhythmicity in relative abundance compared with only 13% (91/717) in naïve IEC-*Bmal1*^−/−^ mice, a relative decrease of over 50% (**Figure 4G**). Given the comparable temporal caloric intake between genotypes, this suggests the host is capable of driving microbiome rhythmicity through IEC-specific *Bmal1*. In mice lacking *Bmal1* in IECs, there was a notable reduction in the proportion of rhythmic OTUs from the Firmicutes phyla, important for host immunity and also reduced in patients with IBD ^35^(**Figure S3D**). IEC-*Bmal1*^−/−^ mice lost rhythmicity of key genera including *Lactobacillus* and *Oscillibacter* (**Figure S3E**), associated with a healthy microbiome ^36^. The majority of genera that lost rhythms in IEC-*Bmal1*^−/−^ mice peaked during the active dark phase (ZT16) in *Bmal1*^flox^ mice (**Figure S3E**).

We additionally set out to explore the effects of localised inflammation on microbial rhythmicity. Microbiome diversity is depleted in mice with DSS colitis and people with IBD, however the impact of these inflammatory conditions on microbiome rhythmicity has not previously been considered. Our analysis revealed a significant reduction in the proportion of rhythmic OTUs (27% vs 14%, p < 0.001) when comparing Day 4 of DSS with the naïve state (**Figure S4A**). Specific genera that lost rhythmicity include *Lactobacillus*, *Bacteroides* and *Roseburia*, all implicated as important for a healthy microbiome. Genera that gained rhythmicity in DSS colitis included *Staphylococcus* and *Ureaplasma*, potential pathogens (**Figure S4B**). In addition, rhythmicity in alpha diversity was lost in mice with DSS colitis (**Figure S4C**). This shows for the first time that DSS colitis dampens rhythmicity in microbiome composition and diversity.

### IEC-specific *Bmal1* deletion does not affect severity of acute DSS colitis

IEC-*Bmal1*^−/−^ and *Bmal1*^flox^ mice treated with 2.5% DSS for 7 days developed colitis, but there were no significant genotype differences in severity (**Figure 5A-5D**). Quantification of circulating inflammatory markers in paired serum samples collected pre-treatment and following 7 days of DSS (**Table S2**) revealed no genotype difference under basal conditions, but DSS induced elevated G-CSF, IL-6 and CXCL1 (**Figure 5E**). The treatment-induced 40-fold increase in serum IL-6 in *Bmal1*^flox^ mice was reduced by over 50% in IEC-*Bmal1*^−/−^ mice, whilst serum IL-1β levels were significantly increased only in IEC-*Bmal1*^−/−^ mice, suggesting IEC-specific *Bmal1* deletion may have subtle effects on local inflammatory processes resulting in an altered profile of circulating disease markers.

**Figure 5:**
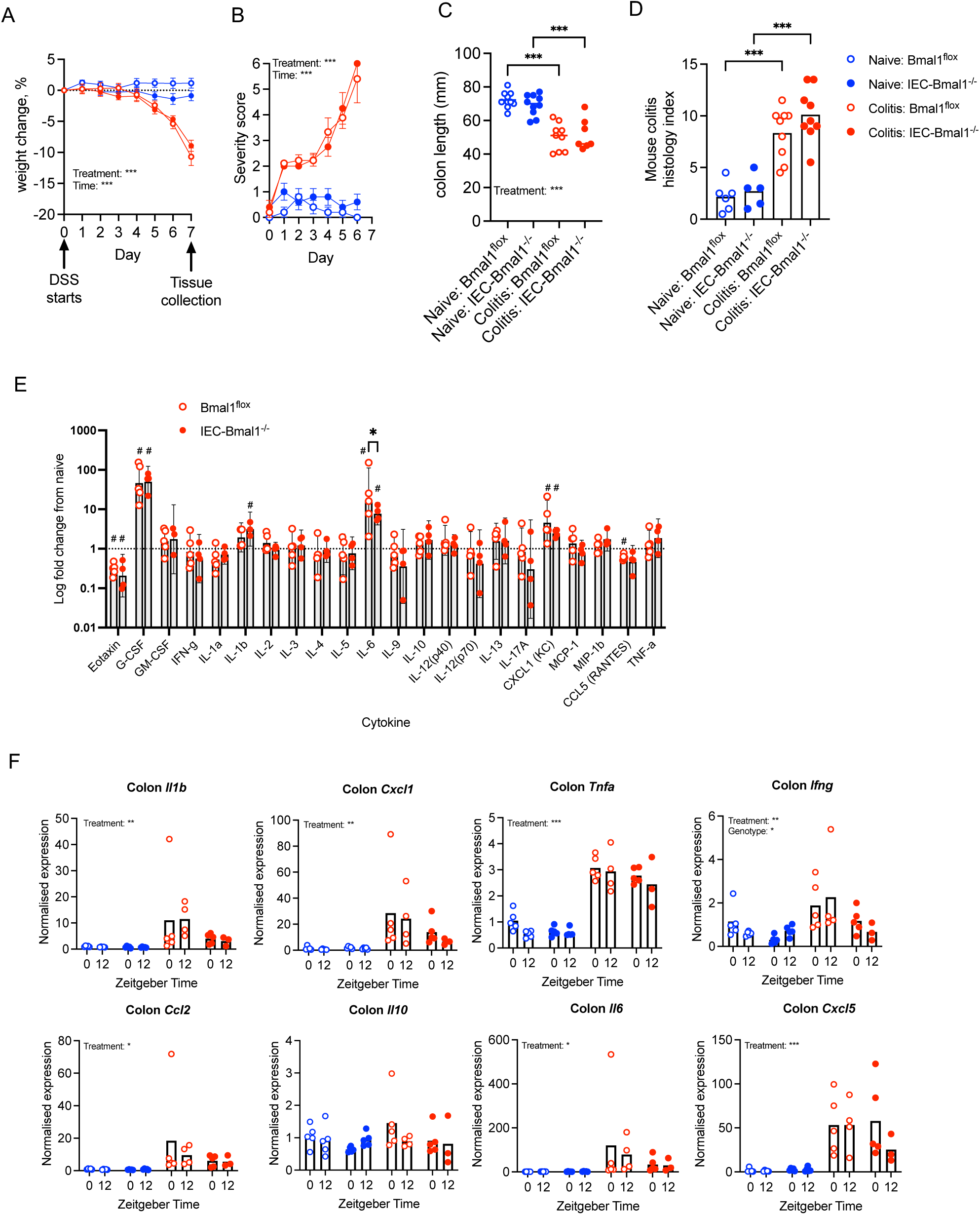
IEC-specific Bmal1 deletion does not affect severity of acute DSS colitis: **(A)** Percentage weight change of mice exposed to water or 2.5% DSS, normalised to day 0 weight, n=8-10/treatment/genotype. **(B)** Daily severity score, n=10/treatment/genotype. **(C)** Colon length measured from distal colon to anus, n=8-10/genotype/treatment. **(D)** Mouse colitis histology index scored on mid-colon samples at day 7, n=5-9/genotype/treatment. **(E)** Log fold change in paired serum cytokine concentration quantified by bio-plex immunoassay pre– and post-DSS. **(F)** qPCR data from colon samples collected at day 7, represented as fold change in ΔΔCt values, using naïve zeitgeber time (ZT)0 as referent population and *Bactin* as housekeeping gene. n=3-5/treatment/genotype/timepoint. Statistics: (A) Three-way ANOVA with multiple comparisons (Tukey). (B) Mixed effects analysis. (C, D) One-way ANOVA with multiple comparisons (Šídák) (E) Two-way ANOVA with multiple comparisons (Šídák)(Asterisk). # represents significant 95% confidence interval (does not include 1.0). (F) Three-way ANOVA with multiple comparisons (Šídák).

To address the impact of *Bmal1* deletion on local inflammatory responses, gene expression of cytokines and chemokines was quantified in colonic samples harvested at ZT0 and ZT12. DSS treatment induced elevated local expression of *Il1b*, C*xcl1*, *Tnfa*, *Ifng*, *Ccl2, Il6* and *Cxcl5* (**Figure 5F**). In contrast to data from C57Bl/6 wildtype mice (**Figure 1J**), *Bmal1*^flox^ animals did not show diurnal variation in gene expression. This likely reflects inter-animal variability in response to this model and limits our ability to conclude on whether local transcription of inflammatory genes does vary by time of day. Only *Ifng* showed a genotype effect, but this did not reach statistical significance in post hoc tests (**Figure 5F**).

### Tregs exhibit treatment-dependent and tissue-dependent rhythmicity

Next, we sought to understand the potential leukocyte subsets contributing to *de novo* rhythmicity in numbers of LP leukocytes from DSS-treated animals. In wildtype mice treated with DSS, numbers of LP T cells, LP CD4^+^ T cells and LP Tregs were robustly rhythmic within the inflamed LP, peaking at ZT12 (**Figure 6A**). The phenomenon of rhythmic Treg numbers under inflammatory conditions is independent of the IEC clock, as further studies in a separate cohort of animals (**Figure 6B**) revealed persistence of diurnal variation in LP Treg numbers in DSS-treated IEC-*Bmal1*^−/−^ mice. Of the other LP leukocyte subsets examined, rhythmicity was not detected in number of neutrophils, monocytes or B cells, but was observed in number of macrophages, conventional dendritic cells and CD8^+^ T cells (**Figure S5A**). Intestinal Tregs are predominantly derived from the thymus or periphery. In mice, it is suggested that thymic Tregs may be distinguished by expression of transcription factor Helios ^37^, although we note that Helios^+^ Tregs can be derived from CD4^+^ conventional T cells under certain contexts ^38^. In *Bmal1*^flox^ mice, following DSS treatment, the number of Helios^+^ Tregs (**Figure 6C**) and the proportion of CD4^+^ T cells (**Figure 6C**) that were Helios^+^ increased at ZT12 in line with total Treg numbers. However, the proportion of Tregs that were Helios^+^ remained unchanged between ZT0 and ZT12 (**Figure 6C**), suggesting an influx of thymic Tregs was unlikely to be driving the peak of LP Tregs at ZT12.

**Figure 6.**
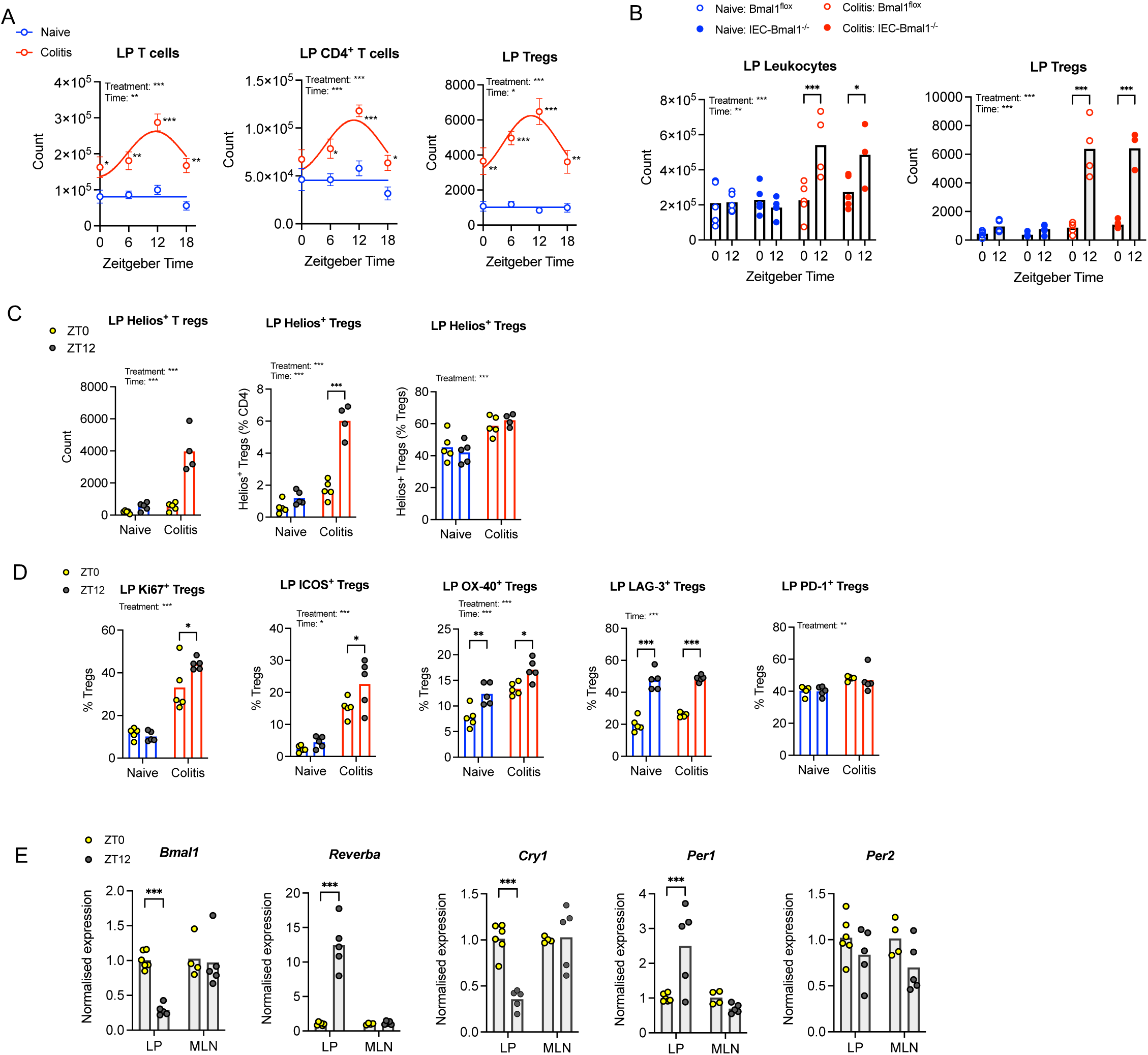
Regulatory T cells exhibit treatment-dependent and tissue-dependent rhythmicity: **(A)** Number of colonic lamina propria (LP) T cells (live CD45^+^CD3^+^), LP CD4^+^ T cells (live CD45^+^CD3^+^CD4^+^) and LP Tregs (live CD45^+^CD3^+^CD4^+^FoxP3^+^) determined by flow cytometry in wildtype mice (n=5/timepoint/treatment/genotype). **(B)** Number of colonic LP leukocytes (live CD45^+^) and LP Tregs (live CD45^+^CD3^+^CD4^+^FoxP3^+^) in IEC-*Bmal1*^−/−^ and *Bmal1*^flox^ mice determined by flow cytometry. N=3-5/timepoint/treatment/genotype. **(C)** Number and proportion of colonic LP Helios^+^ Tregs (live CD45^+^CD3^+^CD4^+^FoxP3^+^Helios^+^), n=4-5/treatment/timepoint. **(D)** Markers of function and proliferation on colonic lamina propria Tregs determined by flow cytometry, n= 5/timepoint/treatment. **(E)** Clock gene expression in Tregs isolated from LP and mesenteric lymph nodes (MLN) of naïve DEREG mice (n=4-6/timepoint), represented as fold change in ΔΔCt values, using naïve zeitgeber time (ZT)0 LP group as referent population and *Gapdh* as housekeeping gene. Statistics: (A, C, D, E) Two-way ANOVA with multiple comparisons (Šídák). (B) Three-way ANOVA with multiple comparisons (Šídák).

To further explore factors driving diurnal Treg numbers, wildtype mice were treated with DSS and LP Treg functional markers assessed by flow cytometry. DSS treatment significantly increased the proportion of Tregs expressing the proliferation marker Ki-67 and ICOS, OX-40 and PD-1 (**Figure 6D**), shown previously to be immunosuppressive ^39^. In naïve animals, LP Tregs exhibited diurnal variation in expression of OX40 and LAG3 which persisted in DSS (**Figure 6D**). Further, in DSS-treated mice, *de novo* diurnal rhythms were observed in the proportion of Tregs expressing Ki-67 and ICOS, peaking at ZT12 (**Figure 6D**). Together, this suggests that within the inflamed colon, LP Treg numbers may peak at ZT12 due to increased intestinal proliferation and the Tregs present at ZT12 are more active.

Previous work in our group demonstrated lack of a functional clock in Tregs isolated from the spleen or inguinal lymph nodes of naïve mice ^40^. However, recent studies have highlighted a functional clock in visceral fat Tregs ^41^. To understand the contribution of the core clock on intestinal Treg function, LP and MLN Tregs were isolated from naïve DEREG mice (which express FoxP3^GFP^, enabling fluorescence-assisted cell sorting of Tregs). Core clock gene expression including *Bmal1*, *Reverbα, Cry1* and *Per1* was robustly diurnal in LP Tregs, but this signal was absent in MLN Tregs (**Figure 6E**). These data support the notion that Treg clock function is tissue-dependent, perhaps driven by factors in the local microenvironment, and reveal the lamina propria as a second tissue site containing Tregs with functional clocks.

## Discussion

This study provides insight into the role of the gut clock on immune and metabolic health, shedding new light on the interplay between the circadian timing system and immunity in the setting of gut inflammation. Cell-specific deletion of *Bmal1* demonstrated a critical role for the IEC clock in regulating mucosal immunity and microbiome rhythmicity. However, the core IEC clock did not reveal itself to be a driver of inflammation in our model of acute DSS-induced colitis. Induction of gut inflammation rapidly disrupted the IEC clock and remodelled the intestinal rhythmic landscape. Our study revealed rhythmic accumulation and activity of anti-inflammatory Tregs within the inflamed colonic LP, with peak abundance and activity during the active phase. This rhythmicity persisted in the absence of local IEC clock. Taken together these data cement the importance of circadian timing in gut health and highlight the need to consider time of day when assessing and treating IBD in the clinic.

There is growing evidence that circadian rhythms modulate gut function, influenced by factors including xenobiotics, dietary and microbiota composition ^6,7,42,43^. RNASeq revealed a critical role for the IEC clock in regulating gut physiology. We found 22% of the naïve wildtype colonic transcriptome showed 24h rhythmicity (which falls within the range of data on rhythmic colonic transcripts from prior work ^6,44^). Rhythmic transcripts mapped onto functions including intestinal immune networks and metabolic processes involving oxidative phosphorylation. We acknowledge that 6-hourly sampling may have limited our detection of cycling transcripts in naïve wildtype mice, compared to similar datasets ^10,45^. IEC-specific deletion of *Bmal1* re-modelled the circadian transcriptome and, in agreement with previous work, was associated with loss of rhythmicity in the microbiota ^46^ and altered dietary fat absorption ^9^. This highlights the importance of the IEC as a timekeeping cell in the gut, orchestrating rhythmic processes critical for immune and metabolic health. Disruption of microbiome rhythmicity in naïve IEC-*Bmal1*^−/−^ mice occurred despite persistent diurnal rhythms in food consumption and rest-activity cycles, demonstrating the importance of the host IEC clock for maintaining microbiome rhythmicity. This challenges the well-documented role of food timing as the main microbiome entrainer (reviewed in ^47^). Our prior work demonstrated feeding-derived rhythms in plasma cell IgA production regulate microbial rhythmicity ^48^. Given that IEC-*Bmal1*^−/−^ mice exhibit robust 24h oscillations in IgA secretion ^48^, rhythms in the microbiota are additionally driven by cues other than feeding time, directly or indirectly via the IEC clock. Potential mechanisms may involve innate anti-microbial peptides, immune sensors such as major histocompatibility complexes and toll-like receptors, and IEC barrier integrity, all demonstrated to interact with the microbiome and be under circadian control ^7,8,49,50^. Whilst our data highlight a role for *Bmal1* in regulating colonic *Claudin 8* expression, this did not translate to perturbation of barrier integrity (as assessed by faecal albumin). Further work is required to determine the relative contribution of host and diet on microbe rhythmicity. Accumulating evidence highlights the importance of robust rhythmic microbiome structure and function for host health including time of day response to pathogen and protection from metabolic and inflammatory disease, suggesting therapeutic potential of restoring microbial rhythmicity ^7,8,51,52^.

Through use of IEC-specific *Reverbα-luciferase* reporter mice, we demonstrated rapid damping of clock gene rhythms in mice administered DSS. Until now, damped clock rhythms have been demonstrated in well-established colitis, via *ex vivo* assays demonstrating weaker PER2::LUC activity in colons from mice ^53^ and clinical studies which demonstrate altered clockwork machinery within inflamed colonic tissue of IBD patients ^13,54^. REV-ERB*α* is recognised to be to be highly sensitive to inflammatory mediators with the protein undergoing rapid proteasomal degradation in response to inflammatory cytokines ^19^. Furthermore, it is established that *Reverbα* transcript levels are reduced during chronic inflammatory insult ^16,40,55,56^, but mechanisms underpinning this are yet to be addressed and consequences of protein degradation on transcript levels not explored. We now add the dimension of time to our understanding of the influence of inflammatory disease on clock components. The core clock responded to inflammatory insult before clinical manifestations were detectable (weight loss or increased severity score). In a clinical context, these data might indicate early loss of rhythmicity in gut function during acute IBD flares, which may have multiple adverse downstream effects, by removing the rhythmic regulation of inflammatory processes and host-microbiome interactions. Thus, thought should be given to the possible benefits of re-instating gut rhythmicity in the setting of colitis.

Despite damping of the local tissue clock in DSS, we observed the rhythmic accumulation and activity of Tregs within the inflamed gut during the active (feeding) phase. Tregs play a critical role in maintaining the intestinal environment under homeostatic conditions, but also suppress inflammation in the setting of colitis ^57^. This anti-inflammatory action occurs via a multitude of mechanisms, including regulation of aberrant T cell activation, tolerance towards luminal antigens, and tissue repair. We predicted that expansion of the Treg population at night would suppress local inflammation in mice. However, transcriptional analysis of gut tissue was unable to conclusively demonstrate day-night variation in the local inflammatory signal within the inflamed gut. This likely reflects inter-animal variation in our model of colitis, but also the heterogenous nature of the colon samples which were tested. It is possible that Treg accumulation at night is associated with a loss of suppressive capacity and therefore did not correlate with a reduced pro-inflammatory signal, however this is unlikely as the presence of Tregs with stable regulatory function is well reported in models of colitis ^58,59^. Further work is warranted to examine individual cellular responses within the inflamed colon across the 24h day.

IEC-specific *Bmal1* deletion did not impact the acute DSS colitis phenotype. Global *Bmal1*^−/−^ mice demonstrate a more severe phenotype in response to DSS ^53,60^, thus our findings implicate the importance of *Bmal1* in other cell types for restraining colonic inflammation. Indeed, the circadian clockwork machinery within both T cells ^61^ and intra-epithelial regulatory B cells ^60^ is important for regulating gut inflammation. We acknowledge that others have shown augmented responses to DSS in mice lacking *Bmal1* in IECs ^10,11^ with this being partially attributed to altered JAK-STAT signalling and microbial metabolites ^11^. These studies used a different Cre driver, TS4Cre ^11^ or floxed mouse model ^10^, which could explain inconsistency between experiments. Additionally, approaches to DSS administration (dose, treatment length, washout) vary between studies and differences in the commensal microbiota between institutes may contribute to variation in responses ^62^.

Robust diurnal oscillations of core clock genes were evident in Tregs isolated from the LP of naïve animals, but not from within MLNs. This aligns with work from our group ^40^ and others ^41^ showing the Treg clock is ticking in non-lymphoid tissues such as visceral adipose, but not lymphoid tissues (spleen and inguinal lymph node). Intestinal Tregs are a unique population, demonstrating a preference for gut-specific trafficking, unlike other non-lymphoid Tregs, which are tissue-agnostic ^63^. It remains to be seen when the intestinal Treg clock starts ticking, but we hypothesise that this may be a response to as yet unidentified tissue-specific signals, which could include inflammatory molecules or microbial-derived signals. We were unable to assess clock status in LP Tregs isolated from mice with colitis, thus cannot determine whether the local inflammatory environment damps the Treg clock as it does local tissue clocks. Given the importance of timing in transplant medicine ^64^, and the potential therapeutic role for Tregs in IBD ^65^ it would be pertinent to consider the clock function of Tregs expanded *ex vivo* and timing of autologous transplant.

In addition to total abundance, Treg suppressive activity was enhanced at the start of the active phase, with higher expression of immunosuppressive molecules including ICOS and LAG3, which has recently been shown to regulate treg metabolism and inflammatory response ^66^. This could be driven by the intrinsic clockwork machinery (if the environment is permissive), or alternatively by daily variation in extrinsic cues including endogenous glucocorticoids, or microbial signals. Glucocorticoids stabilise Treg fate and function in colitis, with Tregs lacking the glucocorticoid receptor failing to suppress disease in the T cell transfer model of colitis ^67^. In support, prior work has demonstrated the importance of daily oscillations in endogenous glucocorticoids in directing rhythmic T cell function ^68^. Further work is warranted exploring the impact of daily variation in glucocorticoid availability on Treg function. This has clinical relevance given that therapeutic glucocorticoids are regularly given (in the morning) to people with acute flares of IBD. Alternatively, data are emerging of bacteria promoting peripheral colonic Treg generation in mice and humans ^69,70^, suggesting microbiome rhythmicity is a candidate extrinsic cue directing daily variation in Treg function. In support, diurnal rhythmicity in levels of glycitein (derived from bacterial breakdown of dietary flavonoids) contributes to daily variation in inflammatory arthritis via actions on pro-inflammatory cells ^52^. It is interesting to speculate around the role of the microbiota in driving rhythmicity in Treg function, but given our discovery that numbers of oscillating bacteria were significantly blunted in colitis, it is important to address whether daily variation in immune modulatory microbial metabolites persist under chronic inflammatory conditions. Further exploration of diurnal variation in these and other extrinsic cues in the setting of colitis and their impact on Treg function may provide new insight into clock regulation on anti-inflammatory mechanisms.

Robust and effective circadian rhythms are integral for immune and metabolic function. Our study further couples the importance of functioning circadian clocks to intestinal health and the impact of colonic inflammation on modulation of circadian processes. The impact of this relationship is multifaceted, and we demonstrate that DSS colitis disrupts existing rhythmic processes, such as IEC core clock expression and microbiome composition, but also that DSS colitis generates *de novo* rhythmicity in processes such as LP Treg abundance and function. Together, this suggests that inflammation perturbs core clock function vital to maintain intestinal homeostasis. Given the increasing prevalence of IBD, the growing awareness of the impact of chrono-disruption on the microbiome and gut health, and the rapid development of both Treg and microbiome-based therapeutics, these findings may help to facilitate the incorporation of circadian logic into therapeutic approaches for treating IBD.

## Acknowledgements

We thank I-Hsuan Lin (Genomic Technologies Core Facility, University of Manchester) for providing support with RNA-sequencing analysis. We thank Gareth Howell (Flow Cytometry Core Facility) for providing support with cell sorting. 16s sequencing data generation and analysis were carried out by the Centre for Genomic Research, which is based at the University of Liverpool. We also thank and acknowledge the University of Manchester Biological Services Facility for animal care. TDB is an NIHR Academic Clinical Lecturer funded by Health Education England (HEE)/NIHR. The views expressed are those of the author(s) and not necessarily those of the NIHR, NHS or the UK Department of Health and Social Care. TDB was funded by a Medical Research Council Clinical Research Training Fellowship during this work (MR/S02199X/1). Work in the Gibbs lab is funded by Versus Arthritis (22625). PD was supported by a Medical Research Council Programme Grant (MR/P023576/1).

## Author Contributions

**TDB**: Funding acquisition, conceptualisation, methodology, formal analysis, investigation, writing – original draft; **PD:** methodology, investigation, formal analysis, writing – review and editing; **SHD:** methodology, investigation; **ALM:** investigation; **DAS:** investigation; **IK:** investigation; **SV:** investigation; **ACW:** methodology, resources; **ADA:** methodology, resources; **DAB:** methodology, resources; **JTM:** Funding acquisition, conceptualisation, writing – review & editing, supervision**; JEG:** Funding acquisition, conceptualisation, methodology, formal analysis, investigation, writing – original draft, supervision

## Supplemental Information

Document S1: Tables S1-S5 and Figures S1-S7

## Resource Availability

### Lead contact

- Requests for further information and resources should be directed to and will be fulfilled by the lead contact, Professor Julie Gibbs.

### Materials availability

- This study did not generate new unique reagents.

### Data and code availability

- RNA sequencing data have been deposited in ArrayExpress (accession code E-MTAB-14341) and are publicly available as of the date of publication.
- This paper does not report original code.
- Any additional information required to reanalyze the data reported in this paper is available from the lead contact upon request.

## STAR Methods

### EXPERIMENTAL MODEL AND STUDY PARTICIPANT DETAILS

#### Animals

Mice were maintained in the University of Manchester Biological Services Facility. Experimental protocols were approved by the University of Manchester Animal Welfare and Ethical Review Body and performed in accordance with the UK Animals (Scientific Procedures) Act 1986. All mice were males, unless stated otherwise. All animals were housed in 12hr light-dark cycles, where Zeitgeber Time 0 (ZT0) represents lights on and ZT12 represents lights off. Animals were fed *ad libitum* standard chow and water, unless specified otherwise. Where documented, light-tight cabinets were used to set timing of lights switching on and off to facilitate tissue collection around the clock whilst maintaining a 12hr light-dark cycle. Mice were acclimatised to cabinets for at least two weeks prior to initiation of experiment.

C57BL/6 mice were purchased from Charles River UK. Homozygous *Βmal1^fl^*^/fl^ mice (JAX, ID: 007668) were crossed with a *Villin*^Cre/+^ line, to produce *Bmal1*^fl/fl^ *Villin*^Cre/+^ (IEC-*Bmal1*^−/−^) offspring and *Bmal1*^fl/fl^ littermate controls (*Bmal1*^flox^) ^71,72^. Hemizygous depletion of regulatory T cell (DEREG) mice with a green fluorescent protein (GFP) fusion protein controlled by FOXP3 promotor/enhancer regions were bred with wildtype counterparts to generate heterozygous offspring and wildtype littermate controls ^73^.

#### Generation of Nr1d1:Stop^fl/fl^Luc mice

*Nr1d1*:Stop^fl/fl^Luc mice with an inducible luciferase reporter construct under the control of the *Nr1d1* (*Reverba*) promoter were created through Bacterial Artificial Chromosome (BAC) recombineering. *Nr1d1*:Stop^fl/fl^Luc mice were crossed with a *Villin*^Cre/+^ line ^71^. The transcription start site was rendered Cre-responsive via insertion of a floxed stop sequence, to produce *Nr1d1*:Stop^fl/fl^Luc *Villin*^Cre/+^ (IEC-*Reverbα*^luc^) offspring with *Nr1d1*:Stop^fl/fl^ wildtype littermate controls.

#### Generation of Nr1d1:Stop^fl/fl^Luc mice: BAC selection

BAC RP23-233I2 was purchased from BACPACchori and used for recombineering. For each BAC recombineering step we followed the protocols developed by the lab of Francis Stewart, Dresden ^74^, using the reagents they kindly provided. First, the conditional reporter, LoxP-STOP-LoxP-Luciferase (LSL-Luc) was integrated in place of the coding region of exon 1 of NR1D1, thus maintaining the critical upstream and intronic regulatory regions previously identified ^75^. This construct contains a strong loxP flanked STOP site ^76^ followed by Luciferase-polyA. Next, a PiggyBac ITR recombination cassette was targeted to the BAC vector ^77,78^. Finally, the bystander genes were removed by the removal of promoter sequences and initial exons through integration of bacterial selection markers (**Figure S6**).

#### Generation of Nr1d1:Stop^fl/fl^Luc mice: Reagent preparation for microinjection

The vector p-mPBase (kind gift from Allan Bradley and Jorge Cadinanos) was used as a template for mPBase mRNA synthesis by mMessage mMachine ULTRA (Ambion). The reaction was cleaned using Megaclear kit (Ambion), eluted in sterile injection buffer (10 mM Tris (pH 7.5), 0.1 mM EDTA (pH 8.0), 100 mM NaCl). BAC DNA was maxi-prepped using BAC Nucleobond kit (Macherey Nagel) and sepharose column purified in sterile injection buffer ^79^. Purified DNA was combined with mPBase mRNA to final concentrations 2 ng/μl BAC and 10 ng/μl mRNA and pronuclear microinjected into one-day single cell C57BL6N mouse embryos. Zygotes were cultured overnight and the resulting 2 cell embryos surgically implanted into the oviduct of day 0.5 post-coitum pseudopregnant mice.

#### Generation of Nr1d1:Stop^fl/fl^Luc mice: Genotyping

After birth and weaning genomic DNA was extracted using Sigma REDextract-n-amp tissue pcr kit according to manufacturer’s instructions. Nine potential founders were identified harbouring the Luciferase transgene but lacking the sequences between the PiggyBac ITRs in the BAC vector, thus indicating transposase mediated single copy integration. A single founder was bred forward to establish a colony and Cre-inducible circadian luminescence confirmed.

#### DSS colitis

Mice (age 8-20 weeks) were administered colitis-grade DSS (36000-50000 molecular weight) (MP Bio, cat: 160110) prepared (2-2.5% weight/volume) in standard water via a spouted bottle *ad libitum* for 6-7 days. Preliminary optimisation studies (using 2-5% DSS) demonstrated that our chosen DSS concentrations provided a robust and repeatable state of colonic inflammation, suitable to interrogate our experimental aims. Mice were weighed daily and colon length from distal caecum to anus recorded at tissue collection. A DSS severity score was utilised (maximum score 12), comprising weight loss, stool consistency and rectal bleeding (**Table S3**).

### METHOD DETAILS

#### Tissue collection

Six-hourly tissue collection timepoints of ZT0, ZT6, ZT12, ZT18 were chosen to represent the start and mid-point of the light and dark cycles, whilst providing adequate time for tissue processing between timepoints. Four-hourly collection timepoints of ZT0, ZT4, ZT8, ZT12, ZT16, ZT20 were chosen to profile microbiome data at higher resolution, which required less inter-timepoint tissue processing.

#### Behavioural analysis

For assessment of rhythmic behaviour, female mice (age 13-19 weeks) were singly housed in cages equipped with running wheels. Wheel running activity was monitored under 12:12 light dark (L:D) for 11 days before switching to constant darkness (D:D). Wheel running activity was recorded throughout using ClockLab software. Data was subsequently analysed using Actogram J in FIJI. Food intake was calculated in the same cohort of animals by weighing food in the hopper at the start of the night (lights off) and start of the day (lights on). The cage floor was swept after each assessment to ensure food could only be consumed from the hoppers. *In vivo* body composition was measured by Body Composition Analyser (EchoMRI) calibrated with canola oil.

#### High fat diet and body composition

High fat diet contained 60% energy from fat (DIO Rodent Purified Diet, IPS Ltd) and was provided *ad libitum* to female mice (14-30 weeks) over a 16-week period. *In vivo* body composition was measured by Body Composition Analyser (EchoMRI) calibrated with canola oil.

#### Histology

Mid-colon tissue was fixed in formalin for 24h and transferred to 70% ethanol. Tissues were embedded in wax blocks and 5μm sections were mounted on glass slides with colons orientated for transverse slices. Slides were dewaxed with xylene and rehydrated in an alcohol series. Nuclei were stained with haematoxylin and counterstained with eosin, using standard protocols. The sections were rinsed in water and dehydrated in alcohol, cleared in xylene and mounted with DPX (Sigma Aldrich). Blinded H&E histology sections were imaged and viewed on QuPath ^80^. Sections were graded using a validated scoring system ^81^.

#### Tissue processing

Colons were stripped of fat, cut longitudinally and washed in cold PBS to remove luminal contents. The tissue was then incubated with 2mM EDTA and 1mM dithiothreitol (DTT) in PBS for 15 minutes on ice, before shaking (30 seconds) and passed through a 70μm filter. The filtered fraction contained IECs ^7,33^, which were isolated by centrifugation at 488x*g* for 5 minutes at 4°C. Flow cytometric assessment for EpCAM expression demonstrated >90% IEC purity (**Figure S7A**). The unfiltered remnant colon was incubated in pre-warmed enzyme digestion cocktail containing collagenase V (0.85mg/ml (final concentration); Sigma), collagenase D (1.25mg/ml; Roche), dispase II (1mg/ml; Gibco) and DNase (30μg/ml; Roche) for 25 minutes. Colons were vortexed and passed through a 40μm filter, immediately washed through with ice cold HBSS with 2% FBS and centrifuged at 488x*g* for 5 minutes at 4°C. To isolate cells from mesenteric lymph nodes (MLN), the largest colonic draining MLN was harvested into cold media with 2% FBS. Cells were mechanically dispersed through a 40μm filter and rinsed with media.

#### RNA extraction and analysis

RNA was extracted from colon tissue, IECs or Tregs following standard protocols. Briefly, colon tissue was homogenised in lysing MatrixD tubes (MP biomedical) containing TRIzol (Invitrogen), using a bead mill homogeniser (Fisherbrand). RNA was extracted with chloroform and aqueous phase was precipitated with isopropyl alcohol. Supernatant was washed with 70% ethanol and the RNA pellet was resuspended in nuclease-free water. RNA was purified from lysed IECs or Tregs with the RNeasy mini kit (Qiagen) or Single cell RNA purification kit (Norgen) according to manufacturer’s instructions.

#### qRT-PCR

RNA was quantified (NanoDrop, ThermoFisher Scientific) and a standardised amount (dependent on tissue and cell type) converted to cDNA using the high-capacity RNA to cDNA kit (Applied Biosystems). qPCR was performed using Takyon ROX Probe Mastermix dTTP blue (Eurogentec) and a StepOnePlus Real-Time PCR machine (Applied Biosystems). Primers and probes are listed in the **Key Resources Table**.

#### RNAseq

RNA was isolated from colonic tissue of naïve IEC-*Bmal1*^−/−^ mice and *Bmal1*^flox^ controls (age 10-20 weeks) at ZT0, ZT6, ZT12 and ZT18. Sequencing library preparation with TruSeq Stranded mRNA assay (Illumina) and sequencing on NovaSeq 6000 instrument (Illumina) were performed by the University of Manchester Genomic Technologies Core Facility. Quality control was performed with Fastqc (v0.11.3) and FastqScreen (v0.14.0). Reads were trimmed using BBDuk from BBMap (v38.96) and reads were mapped to the mouse genome (mm39/vM30) using STAR (v2.7.10a). Differential expression analysis was run in R with DESeq2 using an FDR < 0.05 ^82^. Differential rhythmicity analysis was performed using compareRhythms R package ^27^. Pathway analysis was performed using the Enrichr R package and web tool ^28–30^. Spline plots were generated from RNAseq data utilising all the genes in that pathway that appeared in the dataset.

#### Fecal albumin ELISA

Fecal albumin concentrations were measured using the Mouse Albumin ELISA Quantitation Set (Bethyl Laboratories), according to manufacturer’s instructions, using supernatant from homogenised fecal pellets diluted to 1mg/ml. Absorbance was measured on microplate reader set to 450nm. A standard curve ranging from 1.23-900ng/ml was generated as a sigmoidal 4-parameter curve fit.

#### Multiplex Immunoassay

23-plex Bio-Plex Pro Mouse Cytokine Group 1 assay (BioRad) was performed on paired serum samples obtained from small volume tail bleeds prior to and during DSS treatment at ZT4. Blood samples were centrifuged (10 minutes, 10 000 x g, 4°C) and serum collected and frozen (–80°C) until analysis. Bio-Plex assays were performed according to manufacturer’s instructions, using the 23-plex Bio-Plex Pro Mouse Cytokine Group 1 assay (BioRad). Briefly, serum samples were diluted 1:4 in sample diluent before incubation with coupled beads alongside duplicate standards and blanks. Detection antibody was added, followed by streptavidin-PE. The plate was read on the Bio-Plex 200 system (BioRad).

#### Flow cytometry

Single cell suspensions were analysed on a NucleoCounter NC-250 (ChemoMetec) for viability and cell count. Cell preparations were incubated with anti-CD16/32 (eBioscience) to block Fc receptors. Cells were washed and incubated with extracellular antibodies and live/dead stain for 30 minutes. In the absence of intracellular staining or FACS sorting, cells were fixed in 3.6% formaldehyde for 20 minutes at room temperature in the dark, washed and resuspended in FACS buffer (PBS with 4% FBS and 1mM EDTA). Intracellular staining was performed with the FoxP3/Transcription Factor Staining Buffer Kit (eBioscience). Antibodies directed against cell surface or intracellular markers are listed in the **Key Resources Table**. Stained cells were run on BD LSR Fortessa (BD Biosciences) with BD FACSDiva (BD Biosciences) software and analysed using FlowJo v10 (FlowJo, LLC). Lamina propria CD4^+^FoxP3^+^ Tregs and CD4^+^FoxP3^-^ T cells from diphtheria-naive DEREG mice (age 12-30 weeks) were sorted on the BD Influx (BD Biosciences), with FoxP3^GFP^ detected by the 488nm laser. DEREG mice were chosen for their ease of detecting and isolating Tregs via FoxP3^GFP^ expression. Gating strategies are shown in **Figure S7B-S7E**.

#### In vivo imaging

For *in vivo* bioluminescence imaging, IEC-*Reverbα*^luc^ mice (male and female, 12-22 weeks) were anaesthetised with isoflurane and abdomens were shaved. 10μl/g of 15mg/ml D-luciferin (Promega) was delivered intraperitoneally. Bioluminescence was detected with the *in vivo* imaging system (IVIS) Lumina III (Perkin Elmer). Luciferin kinetic studies determined peak signal intensity between 15-20 minutes after luciferin administration. Total flux was analysed using Living Image software (Perkin Elmer). Villin expression is known to be most abundant in the intestine, with lower levels of expression found in the kidney and also the placenta ^83^. Organs including intestine, kidney, lung, liver and skin were harvested from IEC-*Reverbα*^luc^ and *Nr1d1*:Stop^fl/fl^ controls, incubated with luciferin for 30 minutes and bioluminescence was recorded on a microplate reader (Promega). This demonstrated bioluminescence predominantly from the intestine. The kidneys emitted detectable bioluminescence (**Figure S2F**), however, given their retroperitoneal anatomy and the supine position of mice during bioluminescence acquisition by IVIS *in vivo*, their relative contribution to bioluminescent signal is likely to be negligible.

#### 16S rRNA sequencing and analysis

Microbial DNA was extracted from fecal pellets collected from IEC-*Bmal1*^−/−^ and *Bmal1*^flox^ mice (aged 8-19 weeks) at select times across the 24h day with the DNeasy PowerSoil Pro Kit (Qiagen), as per manufacturer’s instructions. Pre-amplification of the V4 region of *16S* rRNA was performed using forward primer 5’-**ACACTCTTTCCCTACACGACGCTCTTCCGAT-** CTNNNNNGTGCCAGCMGCCGCGGTAA-3’ (annealing sites in bold) and reverse primer 5’-**GTGACTGGAGTTCAGACGTGTGCTCTTCCGATCT**GGACTACHVGGGTWTCTAAT-3’. Sequencing was performed on the Illumina MiSeq v2 platform (Illumina), generating 250bp paired-end reads. PhiX control v3 library (PhiX) was spiked into samples to balance low base diversity often found in microbiome samples. Quality control was performed as described previously^84^. OTU tables were generated using a pipeline provided by the University of Manchester Bioinformatics Core Facility local Galaxy service. Briefly, VSEARCH clustered OTUs and removed chimeras. The OTU database was mapped to the SILVA (v138) reference database with >97% homology threshold. All samples passed quality checks and had sequence depth >45,000. The OTU table was analysed using R packages *phyloseq, vegan*, *limma* and *ALDEx2*. JTK_CYCLE ^85^ was used to identify rhythmic OTUs with a period of 24 h and an adjusted P value < 0.05.

#### Quantification and statistical analysis

Statistical tests were performed in GraphPad Prism or R. Specific statistical tests and corrections for multiple tests are indicated in figure legends. Further detail is provided in **Table S4.** By default, only significant tests are highlighted. Abbreviations: *, p<0.05; **, p<0.01, ***, p<0.001. Graphs were generated in GraphPad Prism or R.

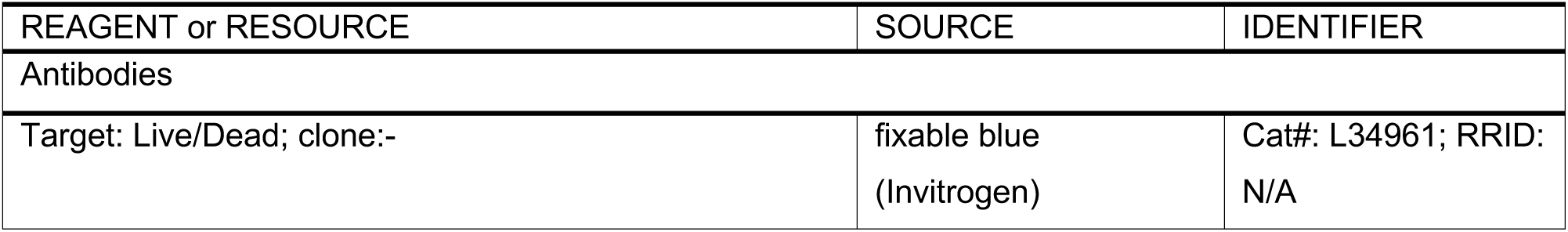
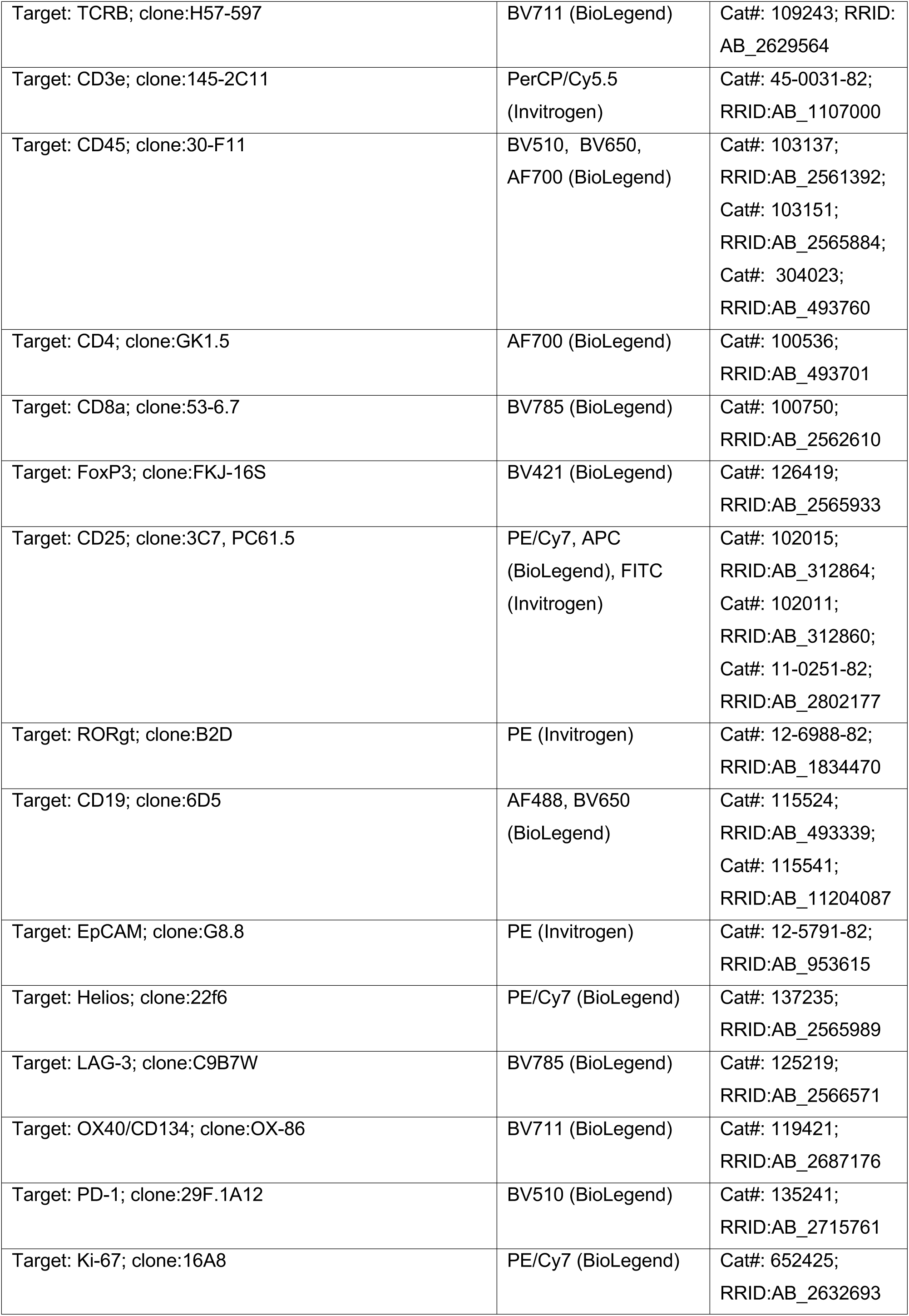
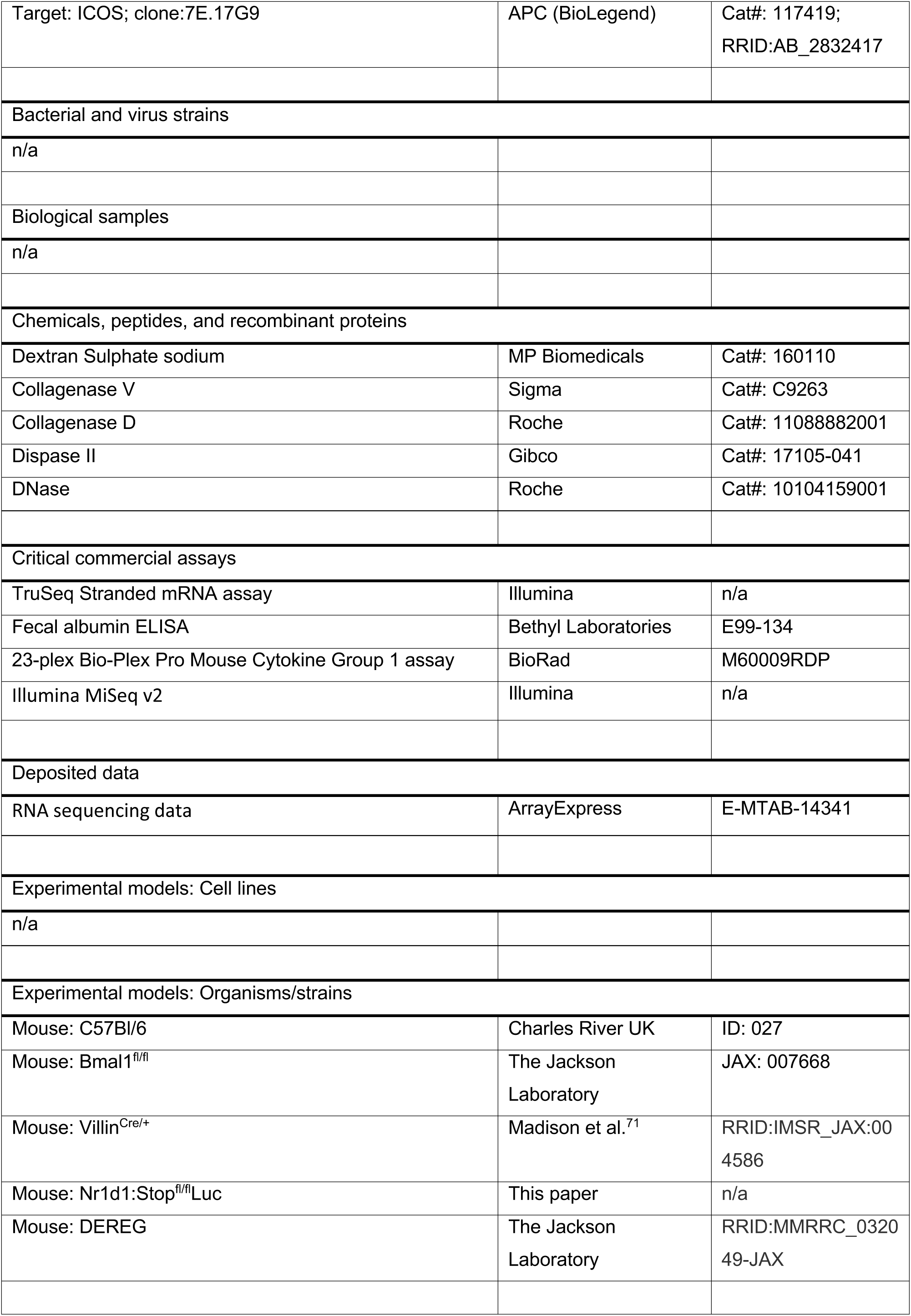
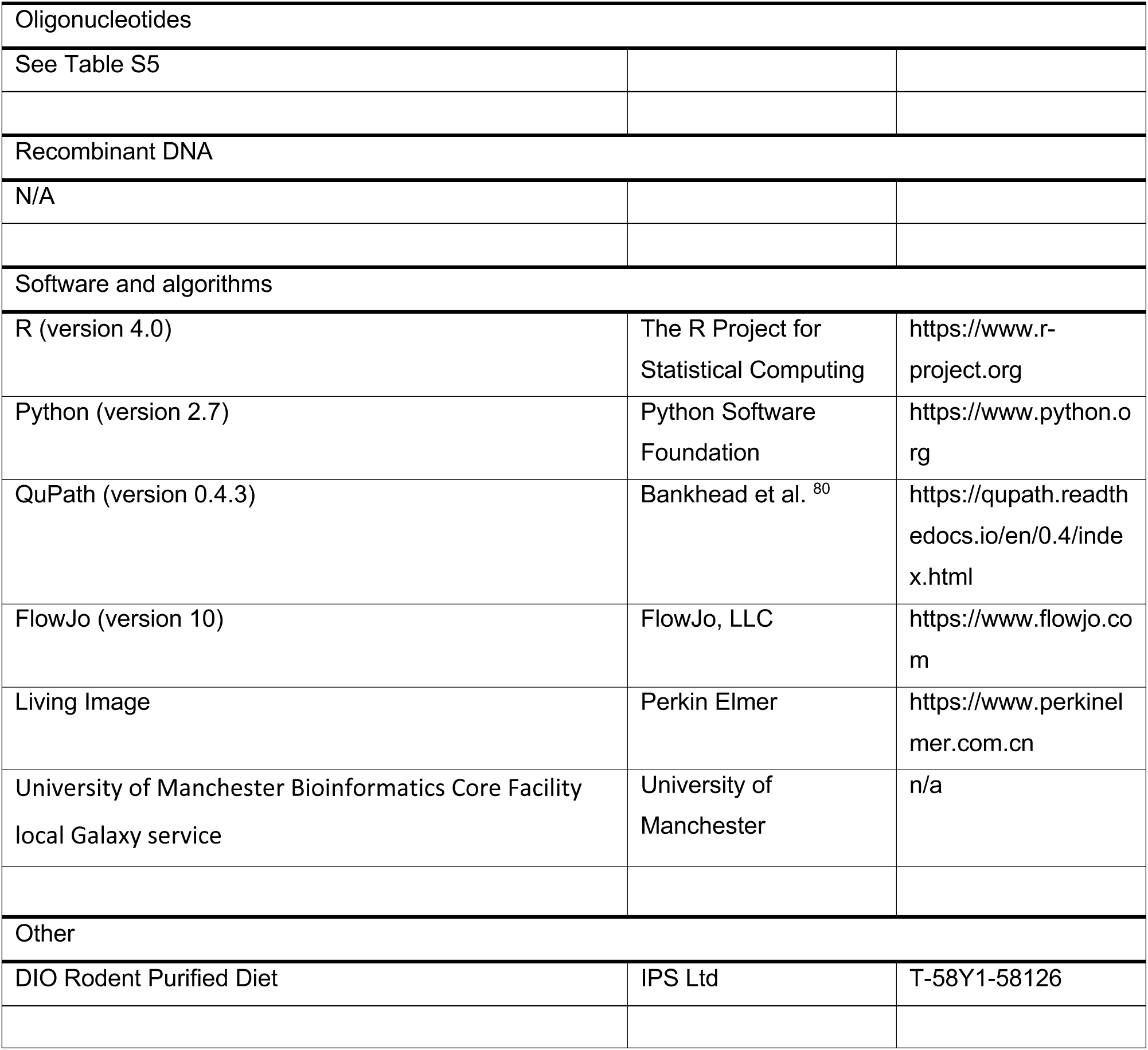
Key resources table.

#### Key resources table

## References

1. de Souza, H.S.P., and Fiocchi, C. (2016). Immunopathogenesis of IBD: current state of the art. Nature Reviews Gastroenterology & Hepatology 13, 13–27. 10.1038/nrgastro.2015.186.

2. Scheiermann, C., Gibbs, J., Ince, L., and Loudon, A. (2018). Clocking in to immunity. Nat Rev Immunol 18, 423–437. 10.1038/s41577-018-0008-4.

3. Sládek, M., Rybová, M., Jindráková, Z., Zemanová, Z., Polidarová, L., Mrnka, L., O’Neill, J., Pácha, J., and Sumová, A. (2007). Insight into the circadian clock within rat colonic epithelial cells. Gastroenterology 133, 1240–1249. 10.1053/j.gastro.2007.05.053.

4. Hoogerwerf, W.A., Hellmich, H.L., Cornélissen, G., Halberg, F., Shahinian, V.B., Bostwick, J., Savidge, T.C., and Cassone, V.M. (2007). Clock gene expression in the murine gastrointestinal tract: endogenous rhythmicity and effects of a feeding regimen. Gastroenterology 133, 1250–1260. 10.1053/j.gastro.2007.07.009.

5. Talbot, J., Hahn, P., Kroehling, L., Nguyen, H., Li, D., and Littman, D.R. (2020). Feeding-dependent VIP neuron–ILC3 circuit regulates the intestinal barrier. Nature 579, 575–580. 10.1038/s41586-020-2039-9.

6. Thaiss, C.A., Levy, M., Korem, T., Dohnalová, L., Shapiro, H., Jaitin, D.A., David, E., Winter, D.R., Gury-BenAri, M., Tatirovsky, E., et al. (2016). Microbiota Diurnal Rhythmicity Programs Host Transcriptome Oscillations. Cell 167, 1495–1510.e1412. 10.1016/j.cell.2016.11.003.

7. Tuganbaev, T., Mor, U., Bashiardes, S., Liwinski, T., Nobs, S.P., Leshem, A., Dori-Bachash, M., Thaiss, C.A., Pinker, E.Y., Ratiner, K., et al. (2020). Diet Diurnally Regulates Small Intestinal Microbiome-Epithelial-Immune Homeostasis and Enteritis. Cell 182, 1441–1459.e1421. 10.1016/j.cell.2020.08.027.

8. Brooks, J.F., II, Behrendt, C.L., Ruhn, K.A., Lee, S., Raj, P., Takahashi, J.S., and Hooper, L.V. (2021). The microbiota coordinates diurnal rhythms in innate immunity with the circadian clock. Cell 184, 4154–4167.e4112. 10.1016/j.cell.2021.07.001.

9. Yu, F., Wang, Z., Zhang, T., Chen, X., Xu, H., Wang, F., Guo, L., Chen, M., Liu, K., and Wu, B. (2021). Deficiency of intestinal Bmal1 prevents obesity induced by high-fat feeding. Nature Communications 12, 5323. 10.1038/s41467-021-25674-5.

10. Niu, Y., Heddes, M., Altaha, B., Birkner, M., Kleigrewe, K., Meng, C., Haller, D., and Kiessling, S. (2024). Targeting the intestinal circadian clock by meal timing ameliorates gastrointestinal inflammation. Cell Mol Immunol. 10.1038/s41423-024-01189-z.

11. Jochum, S.B., Engen, P.A., Shaikh, M., Naqib, A., Wilber, S., Raeisi, S., Zhang, L., Song, S., Sanzo, G., Chouhan, V., et al. (2023). Colonic Epithelial Circadian Disruption Worsens Dextran Sulfate Sodium-Induced Colitis. Inflamm Bowel Dis 29, 444–457. 10.1093/ibd/izac219.

12. Butler, T.D., Mohammed Ali, A., Gibbs, J.E., and McLaughlin, J.T. (2023). Chronotype in Patients With Immune-Mediated Inflammatory Disease: A Systematic Review. J Biol Rhythms 38, 34–43. 10.1177/07487304221131114.

13. Weintraub, Y., Cohen, S., Chapnik, N., Ben-Tov, A., Yerushalmy-Feler, A., Dotan, I., Tauman, R., and Froy, O. (2020). Clock Gene Disruption Is an Initial Manifestation of Inflammatory Bowel Diseases. Clin Gastroenterol Hepatol 18, 115–122.e111. 10.1016/j.cgh.2019.04.013.

14. Huang, M., Wu, Y., Li, Y., Chen, X., Feng, J., Li, Z., Li, J., Chen, J., Lu, Y., and Feng, Y. (2024). Circadian clock-related genome-wide mendelian randomization identifies putatively genes for ulcerative colitis and its comorbidity. BMC Genomics 25, 130. 10.1186/s12864-024-10003-z.

15. Preuss, F., Tang, Y., Laposky, A.D., Arble, D., Keshavarzian, A., and Turek, F.W. (2008). Adverse effects of chronic circadian desynchronization in animals in a “challenging” environment. American Journal of Physiology-Regulatory, Integrative and Comparative Physiology 295, R2034–R2040. 10.1152/ajpregu.00118.2008.

16. Wang, S., Lin, Y., Yuan, X., Li, F., Guo, L., and Wu, B. (2018). REV-ERBα integrates colon clock with experimental colitis through regulation of NF-κB/NLRP3 axis. Nature Communications 9, 4246. 10.1038/s41467-018-06568-5.

17. Amara, J., Saliba, Y., Hajal, J., Smayra, V., Bakhos, J.J., Sayegh, R., and Fares, N. (2019). Circadian Rhythm Disruption Aggravates DSS-Induced Colitis in Mice with Fecal Calprotectin as a Marker of Colitis Severity. Dig Dis Sci 64, 3122–3133. 10.1007/s10620-019-05675-7.

18. Zhang, Z., Li, W., Han, X., Tian, D., Yan, W., Liu, M., and Cao, L. (2024). Circadian rhythm disruption-mediated downregulation of Bmal1 exacerbates DSS-induced colitis by impairing intestinal barrier. Front Immunol 15, 1402395. 10.3389/fimmu.2024.1402395.

19. Pariollaud, M., Gibbs, J.E., Hopwood, T.W., Brown, S., Begley, N., Vonslow, R., Poolman, T., Guo, B., Saer, B., Jones, D.H., et al. (2018). Circadian clock component REV-ERBα controls homeostatic regulation of pulmonary inflammation. The Journal of Clinical Investigation 128, 2281–2296. 10.1172/JCI93910.

20. Downton, P., Sanna, F., Maidstone, R., Poolman, T.M., Hayter, E.A., Dickson, S.H., Ciccone, N.A., Early, J.O., Adamson, A., Spiller, D.G., et al. (2022). Chronic inflammatory arthritis drives systemic changes in circadian energy metabolism. Proc Natl Acad Sci U S A 119, e2112781119. 10.1073/pnas.2112781119.

21. Simpkins, D.A., Downton, P., Gray, K.J., Dickson Suzanna H., Maidstone, R.J., Konkel, J.E., Hepworth Matthew R., Ray, D.W., Bechtold, D.A., and Gibbs, J.E. (2023). Consequences of collagen induced inflammatory arthritis on circadian regulation of the gut microbiome. The FASEB Journal 37, e22704. 10.1096/Ü.202201728R.

22. Liu, A.C., Tran, H.G., Zhang, E.E., Priest, A.A., Welsh, D.K., and Kay, S.A. (2008). Redundant function of REV-ERBalpha and beta and non-essential role for Bmal1 cycling in transcriptional regulation of intracellular circadian rhythms. PLoS Genet 4, e1000023. 10.1371/journal.pgen.1000023.

23. Kortüm, B., Campregher, C., Lang, M., Khare, V., Pinter, M., Evstatiev, R., Schmid, G., Mittlböck, M., Scharl, T., Kucherlapati, M.H., et al. (2015). Mesalazine and thymoquinone attenuate intestinal tumour development in Msh2^loxP/loxP^ Villin-Cre mice. Gut 64, 1905–1912. 10.1136/gutjnl-2014-307663.

24. Pickard, J.M., Maurice, C.F., Kinnebrew, M.A., Abt, M.C., Schenten, D., Golovkina, T.V., Bogatyrev, S.R., Ismagilov, R.F., Pamer, E.G., Turnbaugh, P.J., and Chervonsky, A.V. (2014). Rapid fucosylation of intestinal epithelium sustains host–commensal symbiosis in sickness. Nature 514, 638–641. 10.1038/nature13823.

25. Zhu, L., Han, J., Li, L., Wang, Y., Li, Y., and Zhang, S. (2019). Claudin Family Participates in the Pathogenesis of Inflammatory Bowel Diseases and Colitis-Associated Colorectal Cancer. Front Immunol 10, 1441. 10.3389/fimmu.2019.01441.

26. Li, L., Huang, S., Wang, H., Chao, K., Ding, L., Feng, R., Qiu, Y., Feng, T., Zhou, G., Hu, J.-F., et al. (2018). Cytokine IL9 Triggers the Pathogenesis of Inflammatory Bowel Disease Through the miR21-CLDN8 Pathway. Inflammatory Bowel Diseases 24, 2211–2223. 10.1093/ibd/izy187.

27. Pelikan, A., Herzel, H., Kramer, A., and Ananthasubramaniam, B. (2022). Venn diagram analysis overestimates the extent of circadian rhythm reprogramming. Febs j 289, 6605–6621. 10.1111/febs.16095.

28. Chen, E.Y., Tan, C.M., Kou, Y., Duan, Q., Wang, Z., Meirelles, G.V., Clark, N.R., and Ma’ayan, A. (2013). Enrichr: interactive and collaborative HTML5 gene list enrichment analysis tool. BMC Bioinformatics 14, 128. 10.1186/1471-2105-14-128.

29. Kuleshov, M.V., Jones, M.R., Rouillard, A.D., Fernandez, N.F., Duan, Q., Wang, Z., Koplev, S., Jenkins, S.L., Jagodnik, K.M., Lachmann, A., et al. (2016). Enrichr: a comprehensive gene set enrichment analysis web server 2016 update. Nucleic Acids Res 44, W90–97. 10.1093/nar/gkw377.

30. Xie, Z., Bailey, A., Kuleshov, M.V., Clarke, D.J.B., Evangelista, J.E., Jenkins, S.L., Lachmann, A., Wojciechowicz, M.L., Kropiwnicki, E., Jagodnik, K.M., et al. (2021). Gene Set Knowledge Discovery with Enrichr. Current Protocols 1, e90. 10.1002/cpz1.90.

31. Heuberger, C.E., Janney, A., Ilott, N., Bertocchi, A., Pott, S., Gu, Y., Pohin, M., Friedrich, M., Mann, E.H., Pearson, C., et al. (2023). MHC class II antigen presentation by intestinal epithelial cells fine-tunes bacteria-reactive CD4 T cell responses. Mucosal Immunol. 10.1016/j.mucimm.2023.05.001.

32. Jamwal, D.R., Laubitz, D., Harrison, C.A., Figliuolo da Paz, V., Cox, C.M., Wong, R., Midura-Kiela, M., Gurney, M.A., Besselsen, D.G., Setty, P., et al. (2020). Intestinal Epithelial Expression of MHCII Determines Severity of Chemical, T-Cell-Induced, and Infectious Colitis in Mice. Gastroenterology 159, 1342–1356.e1346. 10.1053/j.gastro.2020.06.049.

33. Beyaz, S., Chung, C., Mou, H., Bauer-Rowe, K.E., Xifaras, M.E., Ergin, I., Dohnalova, L., Biton, M., Shekhar, K., Eskiocak, O., et al. (2021). Dietary suppression of MHC class II expression in intestinal epithelial cells enhances intestinal tumorigenesis. Cell Stem Cell 28, 1922–1935.e1925. 10.1016/j.stem.2021.08.007.

34. Butler, T.D., and Gibbs, J.E. (2020). Circadian Host-Microbiome Interactions in Immunity. Front Immunol 11, 1783. 10.3389/fimmu.2020.01783.

35. Lee, M., and Chang, E.B. (2021). Inflammatory Bowel Diseases (IBD) and the Microbiome-Searching the Crime Scene for Clues. Gastroenterology 160, 524–537. 10.1053/j.gastro.2020.09.056.

36. Durack, J., and Lynch, S.V. (2019). The gut microbiome: Relationships with disease and opportunities for therapy. J Exp Med 216, 20–40. 10.1084/jem.20180448.

37. Thornton, A.M., Korty, P.E., Tran, D.Q., Wohlfert, E.A., Murray, P.E., Belkaid, Y., and Shevach, E.M. (2010). Expression of Helios, an Ikaros transcription factor family member, differentiates thymic-derived from peripherally induced Foxp3+ T regulatory cells. J Immunol 184, 3433–3441. 10.4049/jimmunol.0904028.

38. Pratama, A., Schnell, A., Mathis, D., and Benoist, C. (2020). Developmental and cellular age direct conversion of CD4+ T cells into RORγ+ or Helios+ colon Treg cells. J Exp Med 217. 10.1084/jem.20190428.

39. Nakanishi, Y., Ikebuchi, R., Chtanova, T., Kusumoto, Y., Okuyama, H., Moriya, T., Honda, T., Kabashima, K., Watanabe, T., Sakai, Y., and Tomura, M. (2018). Regulatory T cells with superior immunosuppressive capacity emigrate from the inflamed colon to draining lymph nodes. Mucosal Immunology 11, 437–448. 10.1038/mi.2017.64.

40. Hand, L.E., Gray, K.J., Dickson, S.H., Simpkins, D.A., Ray, D.W., Konkel, J.E., Hepworth, M.R., and Gibbs, J.E. (2020). Regulatory T cells confer a circadian signature on inflammatory arthritis. Nat Commun 11, 1658. 10.1038/s41467-020-15525-0.

41. Xiao, T., Langston, P.K., Muñoz-Rojas, A.R., Jayewickreme, T., Lazar, M.A., Benoist, C., and Mathis, D. (2022). T regs in visceral adipose tissue up-regulate circadian-clock expression to promote fitness and enforce a diurnal rhythm of lipolysis. Science Immunology 7, eabl7641. doi:10.1126/sciimmunol.abl7641.

42. Dantas Machado, A.C., Brown, S.D., Lingaraju, A., Sivaganesh, V., Martino, C., Chaix, A., Zhao, P., Pinto, A.F.M., Chang, M.W., Richter, R.A., et al. (2022). Diet and feeding pattern modulate diurnal dynamics of the ileal microbiome and transcriptome. Cell Rep 40, 111008. 10.1016/j.celrep.2022.111008.

43. Weger, B.D., Gobet, C., Yeung, J., Martin, E., Jimenez, S., Betrisey, B., Foata, F., Berger, B., Balvay, A., Foussier, A., et al. (2019). The Mouse Microbiome Is Required for Sex-Specific Diurnal Rhythms of Gene Expression and Metabolism. Cell Metab 29, 362–382.e368. 10.1016/j.cmet.2018.09.023.

44. Zhang, R., Lahens, N.F., Ballance, H.I., Hughes, M.E., and Hogenesch, J.B. (2014). A circadian gene expression atlas in mammals: implications for biology and medicine. Proc Natl Acad Sci U S A 111, 16219–16224. 10.1073/pnas.1408886111.

45. Simpkins, D.A., Downton, P., Gray, K.J., Dickson, S.H., Maidstone, R.J., Konkel, J.E., Hepworth, M.R., Ray, D.W., Bechtold, D.A., and Gibbs, J.E. (2023). Consequences of collagen induced inflammatory arthritis on circadian regulation of the gut microbiome. Faseb j 37, e22704. 10.1096/Ü.202201728R.

46. Heddes, M., Altaha, B., Niu, Y., Reitmeier, S., Kleigrewe, K., Haller, D., and Kiessling, S. (2022). The intestinal clock drives the microbiome to maintain gastrointestinal homeostasis. Nat Commun 13, 6068. 10.1038/s41467-022-33609-x.

47. Litichevskiy, L., and Thaiss, C.A. (2022). The Oscillating Gut Microbiome and Its Effects on Host Circadian Biology. Annu Rev Nutr 42, 145–164. 10.1146/annurev-nutr-062320-111321.

48. Penny, H.A., Hodge, S.H., and Hepworth, M.R. (2018). Orchestration of intestinal homeostasis and tolerance by group 3 innate lymphoid cells. Seminars in Immunopathology 40, 357–370. 10.1007/s00281-018-0687-8.

49. Mukherji, A., Kobiita, A., Ye, T., and Chambon, P. (2013). Homeostasis in intestinal epithelium is orchestrated by the circadian clock and microbiota cues transduced by TLRs. Cell 153, 812–827. 10.1016/j.cell.2013.04.020.

50. Kyoko, O.O., Kono, H., Ishimaru, K., Miyake, K., Kubota, T., Ogawa, H., Okumura, K., Shibata, S., and Nakao, A. (2014). Expressions of tight junction proteins Occludin and Claudin-1 are under the circadian control in the mouse large intestine: implications in intestinal permeability and susceptibility to colitis. PLoS One 9, e98016. 10.1371/journal.pone.0098016.

51. Reitmeier, S., Kiessling, S., Clavel, T., List, M., Almeida, E.L., Ghosh, T.S., Neuhaus, K., Grallert, H., Linseisen, J., Skurk, T., et al. (2020). Arrhythmic Gut Microbiome Signatures Predict Risk of Type 2 Diabetes. Cell Host Microbe 28, 258–272.e256. 10.1016/j.chom.2020.06.004.

52. Ma, F., Li, Z., Liu, H., Chen, S., Zheng, S., Zhu, J., Shi, H., Ye, H., Qiu, Z., Gao, L., et al. (2024). Dietary-timing-induced gut microbiota diurnal oscillations modulate inflammatory rhythms in rheumatoid arthritis. Cell Metab. 10.1016/j.cmet.2024.08.007.

53. Taleb, Z., Carmona-Alcocer, V., Stokes, K., Haireek, M., Wang, H., Collins, S.M., Khan, W.I., and Karpowicz, P. (2022). BMAL1 Regulates the Daily Timing of Colitis. Frontiers in Cellular and Infection Microbiology 12. 10.3389/fcimb.2022.773413.

54. Wang, D., Yin, H., Wang, X., Wang, Z., Han, M., He, Q., Chen, J., Xian, H., Zhang, B., Wei, X., et al. (2022). Influence of sleep disruption on inflammatory bowel disease and changes in circadian rhythm genes. Heliyon 8, e11229. 10.1016/j.heliyon.2022.e11229.

55. Vasu, V.T., Cross, C.E., and Gohil, K. (2009). Nr1d1, an important circadian pathway regulatory gene, is suppressed by cigarette smoke in murine lungs. Integr Cancer Ther 8, 321–328. 10.1177/1534735409352027.

56. Downton, P., Dickson, S.H., Ray, D.W., Bechtold, D.A., and Gibbs, J.E. (2024). Fibroblast-like synoviocytes orchestrate daily rhythmic inflammation in arthritis. bioRxiv, 2024.2004.2002.587439. 10.1101/2024.04.02.587439.

57. Whibley, N., Tucci, A., and Powrie, F. (2019). Regulatory T cell adaptation in the intestine and skin. Nature Immunology 20, 386–396. 10.1038/s41590-019-0351-z.

58. Yang, B.H., Hagemann, S., Mamareli, P., Lauer, U., Hoffmann, U., Beckstette, M., Föhse, L., Prinz, I., Pezoldt, J., Suerbaum, S., et al. (2016). Foxp3(+) T cells expressing RORγt represent a stable regulatory T-cell effector lineage with enhanced suppressive capacity during intestinal inflammation. Mucosal Immunol 9, 444–457. 10.1038/mi.2015.74.

59. Boschetti, G., Kanjarawi, R., Bardel, E., Collardeau-Frachon, S., Duclaux-Loras, R., Moro-Sibilot, L., Almeras, T., Flourié, B., Nancey, S., and Kaiserlian, D. (2016). Gut Inflammation in Mice Triggers Proliferation and Function of Mucosal Foxp3+ Regulatory T Cells but Impairs Their Conversion from CD4+ T Cells. Journal of Crohn’s and Colitis 11, 105–117. 10.1093/ecco-jcc/jjw125.

60. Liu, J.-L., Wang, C.-Y., Cheng, T.-Y., Rixiati, Y., Ji, C., Deng, M., Yao, S., Yuan, L.-H., Zhao, Y.-Y., Shen, T., and Li, J.-M. (2021). Circadian Clock Disruption Suppresses PDL1+ Intraepithelial B Cells in Experimental Colitis and Colitis-Associated Colorectal Cancer. Cellular and Molecular Gastroenterology and Hepatology 12, 251–276. 10.1016/j.jcmgh.2021.02.008.

61. Amir, M., Chaudhari, S., Wang, R., Campbell, S., Mosure, S.A., Chopp, L.B., Lu, Q., Shang, J., Pelletier, O.B., He, Y., et al. (2018). REV-ERBα Regulates T(H)17 Cell Development and Autoimmunity. Cell Rep 25, 3733–3749.e3738. 10.1016/j.celrep.2018.11.101.

62. Forster, S.C., Clare, S., Beresford-Jones, B.S., Harcourt, K., Notley, G., Stares, M.D., Kumar, N., Soderholm, A.T., Adoum, A., Wong, H., et al. (2022). Identification of gut microbial species linked with disease variability in a widely used mouse model of colitis. Nature Microbiology 7, 590–599. 10.1038/s41564-022-01094-z.

63. Burton, O.T., Bricard, O., Tareen, S., Gergelits, V., Andrews, S., Biggins, L., Roca, C.P., Whyte, C., Junius, S., Brajic, A., et al. (2024). The tissue-resident regulatory T&#xa0;cell pool is shaped by transient multi-tissue migration and a conserved residency program. Immunity 57, 1586–1602.e1510. 10.1016/j.immuni.2024.05.023.

64. Cunningham, P.S., Maidstone, R., Durrington, H.J., Venkateswaran, R.V., Cypel, M., Keshavjee, S., Gibbs, J.E., Loudon, A.S., Chow, C.W., Ray, D.W., and Blaikley, J.F. (2019). Incidence of primary graft dysfunction after lung transplantation is altered by timing of allograft implantation. Thorax 74, 413–416. 10.1136/thoraxjnl-2018-212021.

65. Clough, J.N., Omer, O.S., Tasker, S., Lord, G.M., and Irving, P.M. (2020). Regulatory T-cell therapy in Crohn’s disease: challenges and advances. Gut 69, 942–952. 10.1136/gutjnl-2019-319850.

66. Kim, D., Kim, G., Yu, R., Lee, J., Kim, S., Gleason, M.R., Qiu, K., Montauti, E., Wang, L.L., Fang, D., et al. (2024). Inhibitory co-receptor Lag3 supports Foxp3(+) regulatory T cell function by restraining Myc-dependent metabolic programming. Immunity 57, 2634–2650.e2635. 10.1016/j.immuni.2024.08.008.

67. Rocamora-Reverte, L., Tuzlak, S., von Raffay, L., Tisch, M., Fiegl, H., Drach, M., Reichardt, H.M., Villunger, A., Tischner, D., and Wiegers, G.J. (2019). Glucocorticoid Receptor-Deficient Foxp3+ Regulatory T Cells Fail to Control Experimental Inflammatory Bowel Disease. Frontiers in Immunology 10. 10.3389/fimmu.2019.00472.

68. Shimba, A., Cui, G., Tani-Ichi, S., Ogawa, M., Abe, S., Okazaki, F., Kitano, S., Miyachi, H., Yamada, H., Hara, T., et al. (2018). Glucocorticoids Drive Diurnal Oscillations in T Cell Distribution and Responses by Inducing Interleukin-7 Receptor and CXCR4. Immunity 48, 286–298.e286. 10.1016/j.immuni.2018.01.004.

69. Atarashi, K., Tanoue, T., Shima, T., Imaoka, A., Kuwahara, T., Momose, Y., Cheng, G., Yamasaki, S., Saito, T., Ohba, Y., et al. (2011). Induction of colonic regulatory T cells by indigenous Clostridium species. Science 331, 337–341. 10.1126/science.1198469.

70. Atarashi, K., Tanoue, T., Oshima, K., Suda, W., Nagano, Y., Nishikawa, H., Fukuda, S., Saito, T., Narushima, S., Hase, K., et al. (2013). Treg induction by a rationally selected mixture of Clostridia strains from the human microbiota. Nature 500, 232–236. 10.1038/nature12331.

71. Madison, B.B., Dunbar, L., Qiao, X.T., Braunstein, K., Braunstein, E., and Gumucio, D.L. (2002). Cis elements of the villin gene control expression in restricted domains of the vertical (crypt) and horizontal (duodenum, cecum) axes of the intestine. J Biol Chem 277, 33275–33283. 10.1074/jbc.M204935200.

72. Storch, K.F., Paz, C., Signorovitch, J., Raviola, E., Pawlyk, B., Li, T., and Weitz, C.J. (2007). Intrinsic circadian clock of the mammalian retina: importance for retinal processing of visual information. Cell 130, 730–741. 10.1016/j.cell.2007.06.045.

73. Lahl, K., Loddenkemper, C., Drouin, C., Freyer, J., Arnason, J., Eberl, G., Hamann, A., Wagner, H., Huehn, J., and Sparwasser, T. (2007). Selective depletion of Foxp3+ regulatory T cells induces a scurfy-like disease. J Exp Med 204, 57–63. 10.1084/jem.20061852.

74. Wang, J., Sarov, M., Rientjes, J., Fu, J., Hollak, H., Kranz, H., Xie, W., Stewart, A.F., and Zhang, Y. (2006). An improved recombineering approach by adding RecA to lambda Red recombination. Mol Biotechnol 32, 43–53. 10.1385/mb:32:1:043.

75. Yamamoto, T., Nakahata, Y., Soma, H., Akashi, M., Mamine, T., and Takumi, T. (2004). Transcriptional oscillation of canonical clock genes in mouse peripheral tissues. BMC Mol Biol 5, 18. 10.1186/1471-2199-5-18.

76. Lakso, M., Sauer, B., Mosinger, B., Jr., Lee, E.J., Manning, R.W., Yu, S.H., Mulder, K.L., and Westphal, H. (1992). Targeted oncogene activation by site-specific recombination in transgenic mice. Proc Natl Acad Sci U S A 89, 6232–6236. 10.1073/pnas.89.14.6232.

77. Rostovskaya, M., Fu, J., Obst, M., Baer, I., Weidlich, S., Wang, H., Smith, A.J., Anastassiadis, K., and Stewart, A.F. (2012). Transposon-mediated BAC transgenesis in human ES cells. Nucleic Acids Res 40, e150. 10.1093/nar/gks643.

78. Rostovskaya, M., Naumann, R., Fu, J., Obst, M., Mueller, D., Stewart, A.F., and Anastassiadis, K. (2013). Transposon mediated BAC transgenesis via pronuclear injection of mouse zygotes. Genesis 51, 135–141. 10.1002/dvg.22362.

79. Yang, X.W., Model, P., and Heintz, N. (1997). Homologous recombination based modification in Esherichia coli and germline transmission in transgenic mice of a bacterial artificial chromsome. Nature Biotechnology 15, 859–865. 10.1038/nbt0997-859.

80. Bankhead, P., Loughrey, M.B., Fernández, J.A., Dombrowski, Y., McArt, D.G., Dunne, P.D., McQuaid, S., Gray, R.T., Murray, L.J., Coleman, H.G., et al. (2017). QuPath: Open source software for digital pathology image analysis. Scientific Reports 7, 16878. 10.1038/s41598-017-17204-5.

81. Koelink, P.J., Wildenberg, M.E., Stitt, L.W., Feagan, B.G., Koldijk, M., van ‘t Wout, A.B., Atreya, R., Vieth, M., Brandse, J.F., Duijst, S., et al. (2018). Development of Reliable, Valid and Responsive Scoring Systems for Endoscopy and Histology in Animal Models for Inflammatory Bowel Disease. J Crohns Colitis 12, 794–803. 10.1093/ecco-jcc/jjy035.

82. Love, M.I., Huber, W., and Anders, S. (2014). Moderated estimation of fold change and dispersion for RNA-seq data with DESeq2. Genome Biology 15, 550. 10.1186/s13059-014-0550-8.

83. Luo, Y., Hitz, B.C., Gabdank, I., Hilton, J.A., Kagda, M.S., Lam, B., Myers, Z., Sud, P., Jou, J., Lin, K., et al. (2020). New developments on the Encyclopedia of DNA Elements (ENCODE) data portal. Nucleic Acids Res 48, D882–d889. 10.1093/nar/gkz1062.

84. Schirmer, M., Ijaz, U.Z., D’Amore, R., Hall, N., Sloan, W.T., and Quince, C. (2015). Insight into biases and sequencing errors for amplicon sequencing with the Illumina MiSeq platform. Nucleic Acids Research 43, e37–e37. 10.1093/nar/gku1341.

85. Hughes, M.E., Hogenesch, J.B., and Kornacker, K. (2010). JTK_CYCLE: an efficient nonparametric algorithm for detecting rhythmic components in genome-scale data sets. J Biol Rhythms 25, 372–380. 10.1177/0748730410379711.

