## Supplementary data for "Colonic inflammation modulates the intestinal circadian landscape"

**This PDF file includes:**

Supplementary Tables 1-5
Supplementary Figures 1-7

**Table S1.** Genes included in pathway analysis

| KEGG 2019 mouse Pathway | Number of genes | Genes in pathway |
|---|---|---|
| IBD (mmu05321) | 54 | foxp3, h2-aa, h2-ab1, h2-dma, h2-dmb1, h2-dmb2, h2-eb1, h2-oa, h2-ob, ifng, ifngr1, ifngr2, il10, il12a, il12b, il12rb1, il12rb2, il13, il17a, il17f, il18, il18r1, il18rap, il1a, il1b, il21, il21r, il22, il23a, il23r, il2rg, il4ra, jun, maf, nfatc1, nfkb1, nod2, rela, rora, rorc, smad2, smad3, stat1, stat3, stat4, stat6, tbx21, tgfb1, tgfb2, tgfb3, tlr2, tlr4, tlr5, tnf |
| Intestinal immune network for IgA production (mmu04672) | 37 | aicda, ccl25, ccl28, ccr10, ccr9, cd28, cd40, cd40lg, cd80, cd86, cxcl12, cxcr4, h2-aa, h2-ab1, h2-dma, h2-dmb1, h2-dmb2, h2-eb1, h2-oa, h2-ob, icos, icosl, il10, il15, il15ra, itga4, itgb7, ltbr, madcam1, map3k14, pigr, tgfb1, tnfrsf13b, tnfrsf13c, tnfrsf17, tnfsf13, tnfsf13b |
| Antigen processing and presentation (mmu04612) | 71 | b2m, calr, canx, cd4, cd74, cd8a, cd8b1, ciita, creb1, ctsb, ctsl, ctss, gm7030, gm8909, h2-aa, h2-ab1, h2-bl, h2-d1, h2-dma, h2-dmb1, h2-dmb2, h2-eb1, h2-k1, h2-m2, h2-m3, h2-m5, h2-m9, h2-oa, h2-ob, h2-q1, h2-q10, h2-q2, h2-q4, h2-q6, h2-q7, h2-t-ps, h2-t10, h2-t22, h2-t23, h2-t24, h2-t3, hsp90aa1, hsp90ab1, hspa1a, hspa1b, hspa1l, hspa2, hspa4, hspa5, hspa8, ifi30, ifng, klrc1, klrc2, klrd1, lgmn, nfya, nfyb, nfyc, pdia3, psme1, psme2, psme2b, psme3, rfx5, rfxank, rfxap, tap1, tap2, tapbp, tnf |
| Tight junction (mmu04530) | 147 | actb, actg1, actn1, actn4, actr2, actr3, actr3b, afdn, amot, amotl1, amotl2, arhgap17, arhgef18, arhgef2, bves, cacna1d, ccnd1, cd1d1, cdc42, cdk4, cftr, cgn, cgnl1, cldn1, cldn10, cldn11, cldn14, cldn15, cldn2, cldn20, cldn22, cldn23, cldn3, cldn4, cldn5, cldn7, cldn8, cldn9, crb3, cttn, dlg1, dlg2, dlg3, epb41l4b, erbb2, ezr, f11r, gata4, hcls1, hspa4, igsf5, itgb1, jam2, jam3, jun, llgl1, llgl2, magi1, map2k7, map3k1, map3k5, mapk10, mapk8, mapk9, marveld2, marveld3, micall2, mpdz, mpp4, msn, myh10, myh11, myh14, myh2, myh3, myh7b, myh9, myl12a, myl12b, myl6, myl6b, myl9, nedd4, nedd4l, nf2, ocln, pard3, pard6a, pard6b, pard6g, patj, pcna, ppp2ca, ppp2cb, ppp2r1a, ppp2r1b, ppp2r2a, ppp2r2b, ppp2r2c, ppp2r2d, prkaa1, prkaa2, prkab1, prkab2, prkaca, prkacb, prkag1, prkag2, prkag3, prkce, prkci, prkcz, rab13, rab8a, rab8b, rac1, rap1a, rap2c, rapgef2, rapgef6, rdx, rhoa, rock1, rock2, runx1, scrib, slc9a3r1, src, stk11, sympk, synpo, tiam1, tjap1, tjp1, tjp2, tjp3, tuba1a, tuba1b, tuba1c, tuba4a, tuba8, tubal3, vasp, was, wasl, whamm, ybx3 |

**Table S2:** Baseline cytokine levels from multiplex immunoassay

| | Bmal1$^{flox}$ | | IEC-Bmal1$^{-/-}$ | | |
|---|---|---|---|---|---|
| **Cytokine** | **Mean** (pg/ml) | **SEM** (pg/ml) | **Mean** (pg/ml) | **SEM** (pg/ml) | **p value** |
| **Eotaxin** | 3444.8 | 309.9 | 3529.4 | 650.8 | 0.90 |
| **G-CSF** | 151.5 | 45.0 | 163.1 | 50.2 | 0.87 |
| **GM-CSF** | 58.0 | 10.5 | 50.0 | 15.0 | 0.67 |
| **IFN-γ** | 62.3 | 32.4 | 34.3 | 7.3 | 0.48 |
| **IL-1α** | 153.2 | 35.0 | 65.3 | 6.2 | 0.06 |
| **IL-1β** | 19.0 | 2.6 | 9.3 | 3.0 | 0.06 |
| **IL-2** | 13.1 | 1.7 | 12.5 | 1.3 | 0.79 |
| **IL-3** | 11.1 | 1.1 | 9.6 | 2.8 | 0.60 |
| **IL-4** | 15.5 | 1.5 | 12.7 | 2.7 | 0.38 |
| **IL-5** | 18.5 | 3.4 | 17.1 | 5.3 | 0.83 |
| **IL-6** | 8.4 | 4.6 | 11.7 | 4.7 | 0.64 |
| **IL-9** | 49.3 | 23.2 | 27.6 | 5.6 | 0.44 |
| **IL-10** | 65.1 | 5.4 | 246.5 | 181.1 | 0.29 |
| **IL-12(p40)** | 1122.2 | 108.4 | 1033.0 | 96.0 | 0.57 |
| **IL-12(p70)** | 208.7 | 32.6 | 219.3 | 52.0 | 0.86 |
| **IL-13** | 77.7 | 15.0 | 52.5 | 9.1 | 0.22 |
| **IL-17A** | 194.6 | 34.0 | 170.1 | 35.2 | 0.63 |
| **CXCL1 (KC)** | 143.4 | 51.6 | 172.0 | 27.0 | 0.66 |
| **MCP-1** | 477.4 | 110.1 | 583.0 | 131.4 | 0.55 |
| **MIP-1α** | 8.4 | 0.8 | 5.9 | 0.8 | 0.06 |
| **MIP-1β** | 61.2 | 4.7 | 46.8 | 3.8 | 0.06 |
| **CCL5 (RANTES)** | 291.2 | 24.2 | 216.7 | 23.3 | 0.07 |
| **TNF-α** | 153.7 | 15.4 | 181.3 | 58.1 | 0.62 |

**Table S3:** DSS colitis scoring system

| Score | Weight loss | Stool consistency | Bleeding |
|---|---|---|---|
| 0 | 0 | Normal | No blood |
| 1 | 5-10% | - | - |
| 2 | 10-15% | Loose | Visible blood on pellet |
| 3 | 15-20% | - | - |
| 4 | >20% | Diarrhoea | Gross bleeding with blood around anus |

**Table S4.** Summary of statistical tests and results

Table S4 is presented in a separate .xls file

**Table S5:** *qPCR* primers and probes

| Gene | Forward primer | Reverse primer | Fam-Tamra probe |
|---|---|---|---|
| *Gapdh* | CAA TGT GTC CGT CGT CGA TCT | GTC CTC AGT GTA GCC CAA GAT G | CGT GCC GCC TGG AGA AAC CTG CC |
| *Rps18* | Proprietary assay (Mm02601777_g1) | | |
| *β-actin* | AGG TCA TCA CTA TTG GCA ACG A | CAC TTC ATG ATG GAA TTG AAT GTA GTT | TGC CAC AGG ATT CCA TAC CCA AGA AGG |
| *Cre* | Proprietary assay (Mr00635245_cn) | | |
| *Bmal1 (exon 8)* | CGT CGG GAC AAA ATG AAC AG | GAA CAG CCA TCC TTA GCA C | TAC CCA CAT GCA ATG CAA TGT CCA GGA A |
| *Nr1d1 (Reverbα)* | Proprietary assay (Mm00520708_m1) | | |
| *Per2* | GCC TTC AGA CTC ATG ATG ACA GA | TTT GTG TGC GTC AGC TTT GG | ACT GCT CAC TAC TGC AGC CGC TCG T |
| *Cry1* | CTGGCGTGG AAGTCATCGT | CTGTCCGCCATT GAGTTCTATG | CGCATTTCACATACACT GTATGACCTGGACA |
| *Il1β* | TCG CTC AGG GTC ACA AGA AA | CCA TCA GAG GCA AGG AGG AA | CAT GGC ACA TTC TGT TCA AAG AGA GCC TG |
| *Il6* | CTA TAC CAC TTC ACA AGT CGG AGG | TGC ACA ACT CTT TTC TCA TTT CC | TTA ATT ACA CAT GTT CTC TGG GAA ATC G |
| *Il10* | Proprietary assay (Mm01288386_m1) | | |
| *Tnfα* | TCT CTT CAA GGG ACA AGG CTG | ATA GCA AAT CGG CTG ACG GT | CCC GAC TAC GTG CTC CTC ACC CA |
| *Ifnγ* | TCA AGT GGC ATA GAT GTG GAA GAA | TGG CTC TGC AGG ATT TTC ATG | TCA CCA TCC TTT TGC CAG TTC CTC CAG |
| *Cxcl1* | CTG CAC CCA AAC CGA AGT | AGC TTC AGG GTC AAG GCA AG | CAC TCA AGA ATG GTC GCG AGG C |
| *Cxcl5* | Proprietary assay (Mm00436451_ g1) | | |
| *Ccl2* | TTC TGG GCC TGC TGT TCA | CCA GCC TAC TCA TTG GGA TCA | CTC AGC CAG ATG CAG TTA ACG CCC C |

### Figure S1

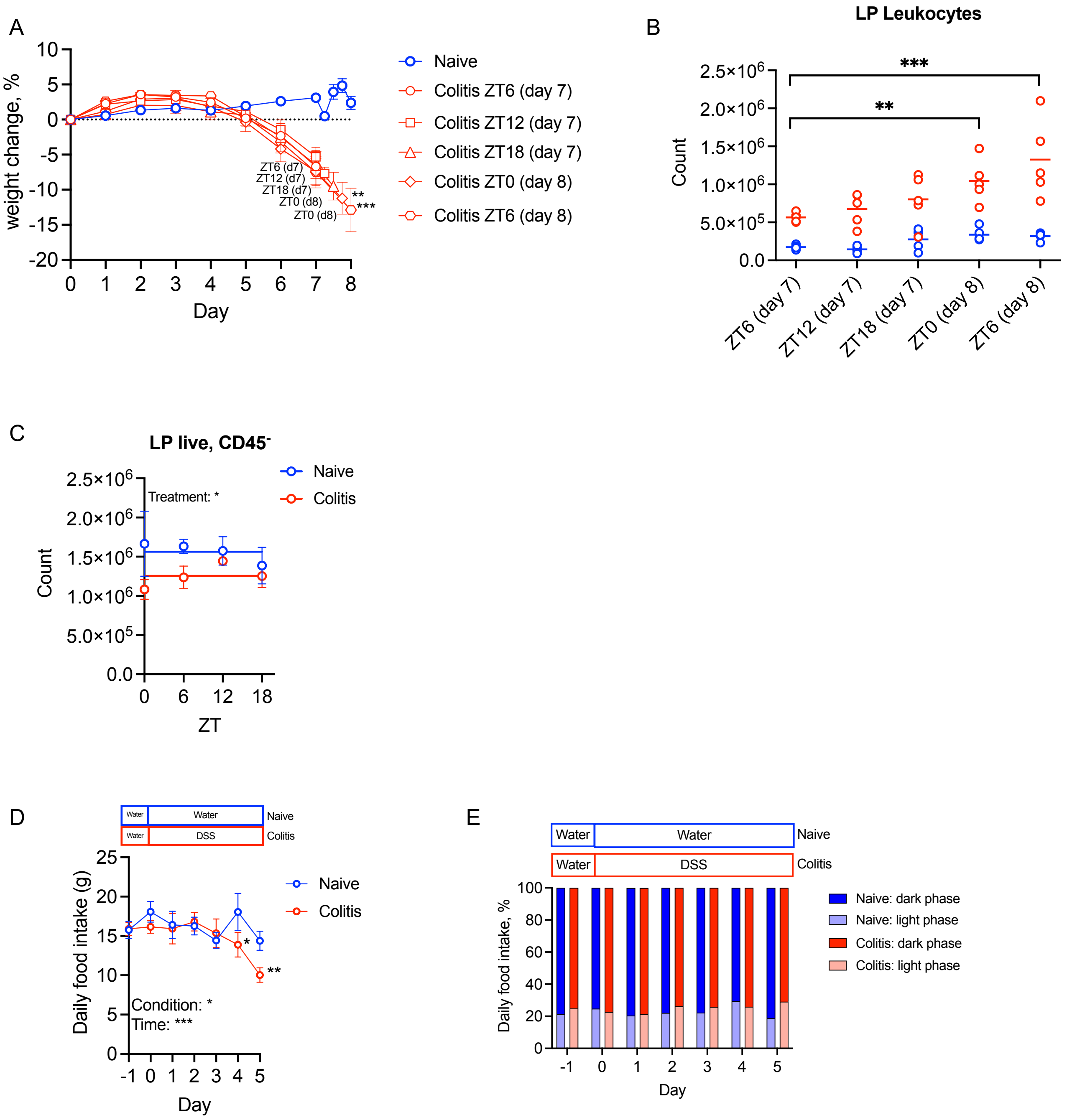

A
weight change, %
Day
Naive
Colitis ZT6 (day 7)
Colitis ZT12 (day 7)
Colitis ZT18 (day 7)
Colitis ZT0 (day 8)
Colitis ZT6 (day 8)
ZT6 (d7)
ZT12 (d7)
ZT18 (d7)
ZT0 (d8)
ZT0 (d8)
**
***
B
LP Leukocytes
Count
***
**
ZT6 (day 7)
ZT12 (day 7)
ZT18 (day 7)
ZT0 (day 8)
ZT6 (day 8)
C
LP live, CD45-
Treatment: *
Count
ZT
Naive
Colitis
D
Water
Water
Naive
Water
DSS
Colitis
Daily food intake (g)
Day
Naive
Colitis
Condition: *
Time: ***
E
Water
Water
Naive
Water
DSS
Colitis
Daily food intake, %
Day
Naive: dark phase
Naive: light phase
Colitis: dark phase
Colitis: light phase

**Figure S1:** **(A)** Percentage weight change of mice exposed to water or 2.5% DSS initiated at the same time, Zeitgeber Time (ZT)4), where ZT0 represents lights on, and ZT12 represents lights off. Data were normalised to day 0 weight (DSS treated, n=5/timepoint; Naïve controls, n=5/timepoint (data pooled into one group). **(B)** Number of lamina propria (LP) leukocytes (live $CD45^{+}$), assessed by flow cytometry, n=5/timepoint/treatment. **(C)** Flow cytometry data showing number of live $CD45^{-}$ cells in the lamina propria across time in naïve and DSS treated (colitis) mice, n=5/treatment/timepoint. **(D)** Daily food intake prior to the initiation of colitis and during DSS treatment (per cage, n=4/cage, 1 cage/treatment) **(E)** Day/night food intake (per cage, n=4/cage, 1 cage/treatment) as a percentage of total daily food intake. Statistics: (A) Two-way ANOVA with multiple comparisons (Šídák). * = compared with colitis ZT6 (day 7). (B) Two-way ANOVA with multiple comparisons (Dunnett) (C) Two-way ANOVA with multiple comparisons (Šídák) and nonlinear regression to compare whether best fit is given by horizontal line or sine wave with nonzero baseline, constraints: wavelength = 24 hours; amplitude > 0. (D) Two-way ANOVA with multiple comparisons (Tukey).

**Figure S2**

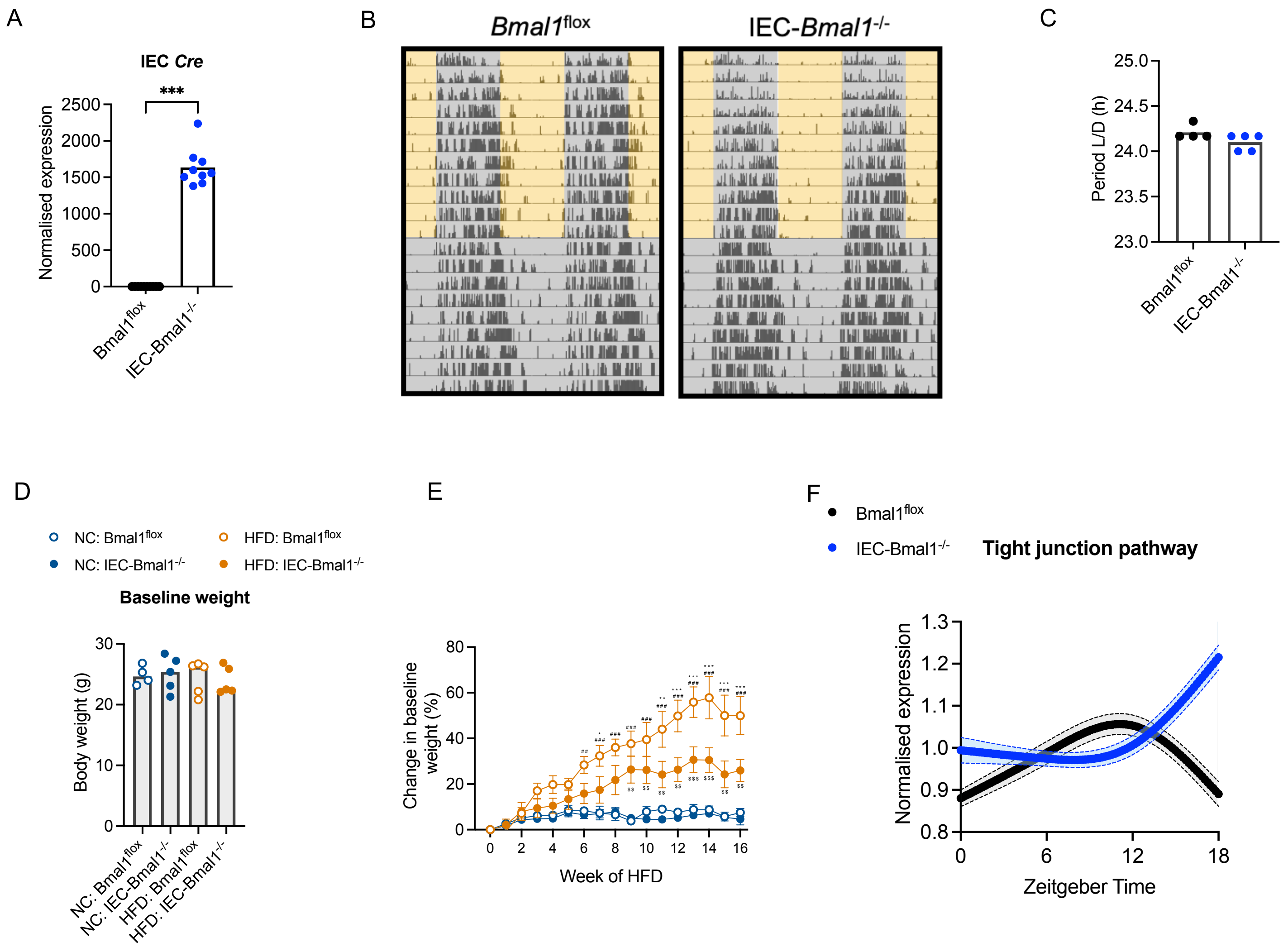
A
IEC Cre
***
Normalised expression
2500
2000
1500
1000
500
0
Bmal1flox
IEC-Bmal1-/-
B
Bmal1flox
IEC-Bmal1-/-
C
Period L/D (h)
25.0
24.5
24.0
23.5
23.0
Bmal1flox
IEC-Bmal1-/-
D
NC: Bmal1flox
NC: IEC-Bmal1-/-
HFD: Bmal1flox
HFD: IEC-Bmal1-/-
Baseline weight
Body weight (g)
30
20
10
0
NC: Bmal1flox
NC: IEC-Bmal1-/-
HFD: Bmal1flox
HFD: IEC-Bmal1-/-
E
Change in baseline weight (%)
80
60
40
20
0
0
2
4
6
8
10
12
14
16
Week of HFD
F
Bmal1flox
IEC-Bmal1-/-
Tight junction pathway
Normalised expression
1.3
1.2
1.1
1.0
0.9
0.8
0
6
12
18
Zeitgeber Time

**Figure S2: (A)** Colonic intestinal epithelial cell (IEC) *Cre* gene expression in naïve mice, represented as fold change in ΔΔCt values, using naïve ZT0 group as referent population and *Gapdh* as housekeeping gene, n=9-10/genotype. **(B)** Representative wheel-running actogram in light/dark (Day 1-11) and constant dark (Day 12-20) conditions. **(C)** Actogram-derived period length during light/dark conditions, n=4-5/genotype. **(D)** Baseline body weight of animals prior to high fat diet (HFD; n=4-5/genotype) or for controls normal chow (NC; n=5/genotype). **(E)** Percentage weight change in mice during 16 week HFD or NC, normalised to baseline weight. * Significance vs. HFD: IEC-*Bmal1*$^{-/-}$; $^{\#}$ Significance vs. NC: *Bmal1*$^{flox}$; $^{\$}$ Significance vs. NC: IEC-*Bmal1*$^{-/-}$. **(F)** Spline plots showing mean normalised expression of all genes from the dataset within the tight junction pathway (mmu04530 147/169), error bars represent 95% confidence intervals. Statistics: (A) unpaired two-tailed t test (C) unpaired two-tailed t test (D) One-way ANOVA (E) Two-way ANOVA with multiple comparisons (Tukey).

### Figure S3

A

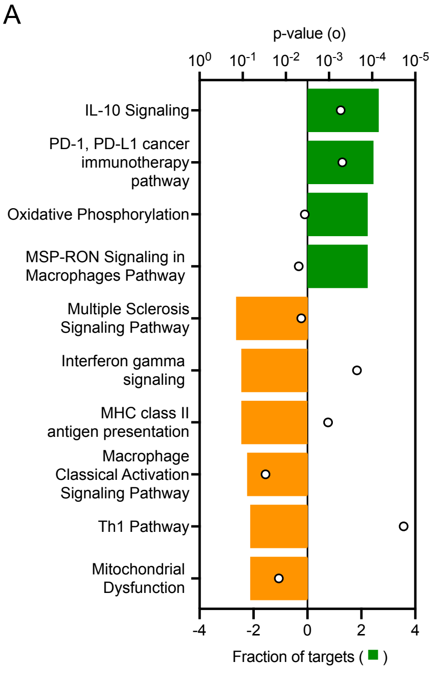


B

| Diseases or Functions Annotation | p-value |
|---|---|
| Chronic colitis | 1.82E-12 |
| Experimental colitis | 3.80E-12 |
| Epithelial neoplasm | 3.99E-11 |
| Non-melanoma solid tumor | 7.61E-11 |
| Carcinoma | 1.19E-10 |
| Endocrine gland tumor | 1.23E-10 |
| Non-hematological solid tumor | 1.89E-10 |
| Endocrine carcinoma | 3.10E-10 |
| Dextran sodium sulfate-induced colitis | 3.94E-10 |
| Nonpituitary endocrine tumour | 4.78E-10 |

C

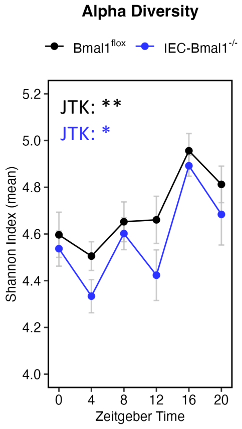


D

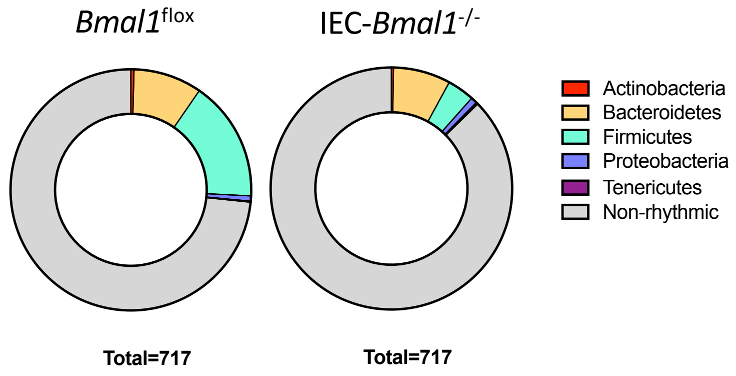


E

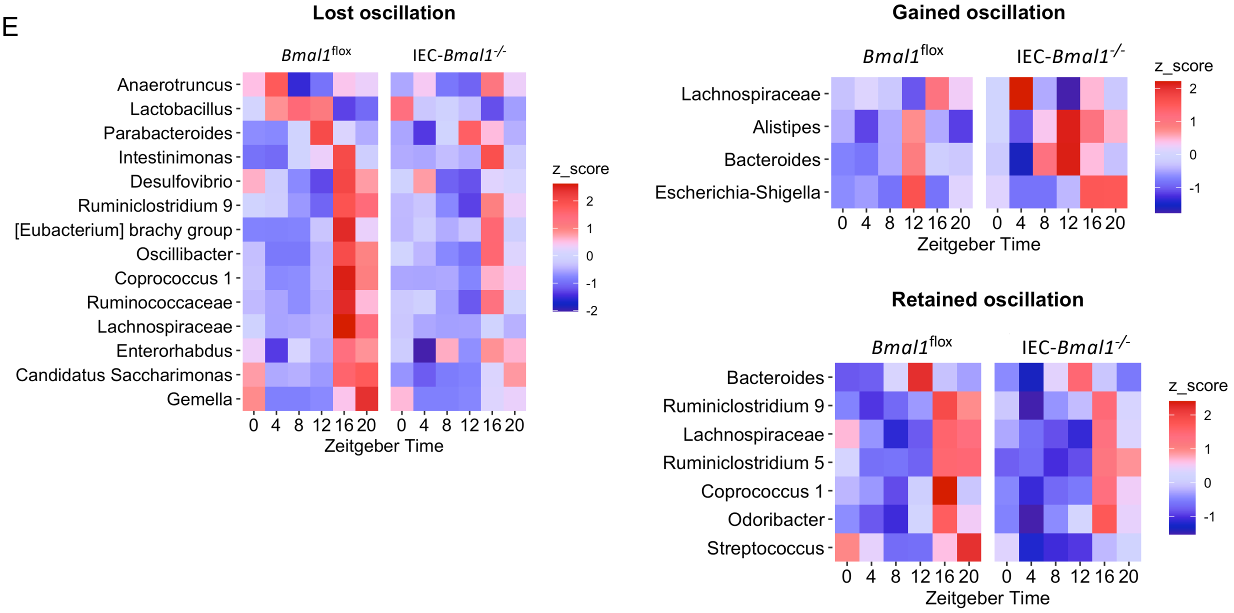

**Figure S3: (A)** Ingenuity pathway analysis of significantly activated (green, z score >2) and inhibited (orange, z score > -2) pathways in colonic transcriptome from IEC-*Bmal1*$^{-/-}$ mice compared to *Bmal1*$^{flox}$. **(B)** A table of the top 10 disease annotations significantly impacted in IEC-*Bmal1*$^{-/-}$ mice compared to *Bmal1*$^{flox}$. **(C)** Alpha diversity within 16S microbiome sequencing (n=30/genotype). **(D)** Rhythmic operational taxonomic unit (OTUs) categorised by phyla. **(E)** Heatmaps of comparative rhythmicity for annotated genera. Statistics: (C and E) Analysis of rhythmicity by JTK_CYCLE.

#### Figure S4

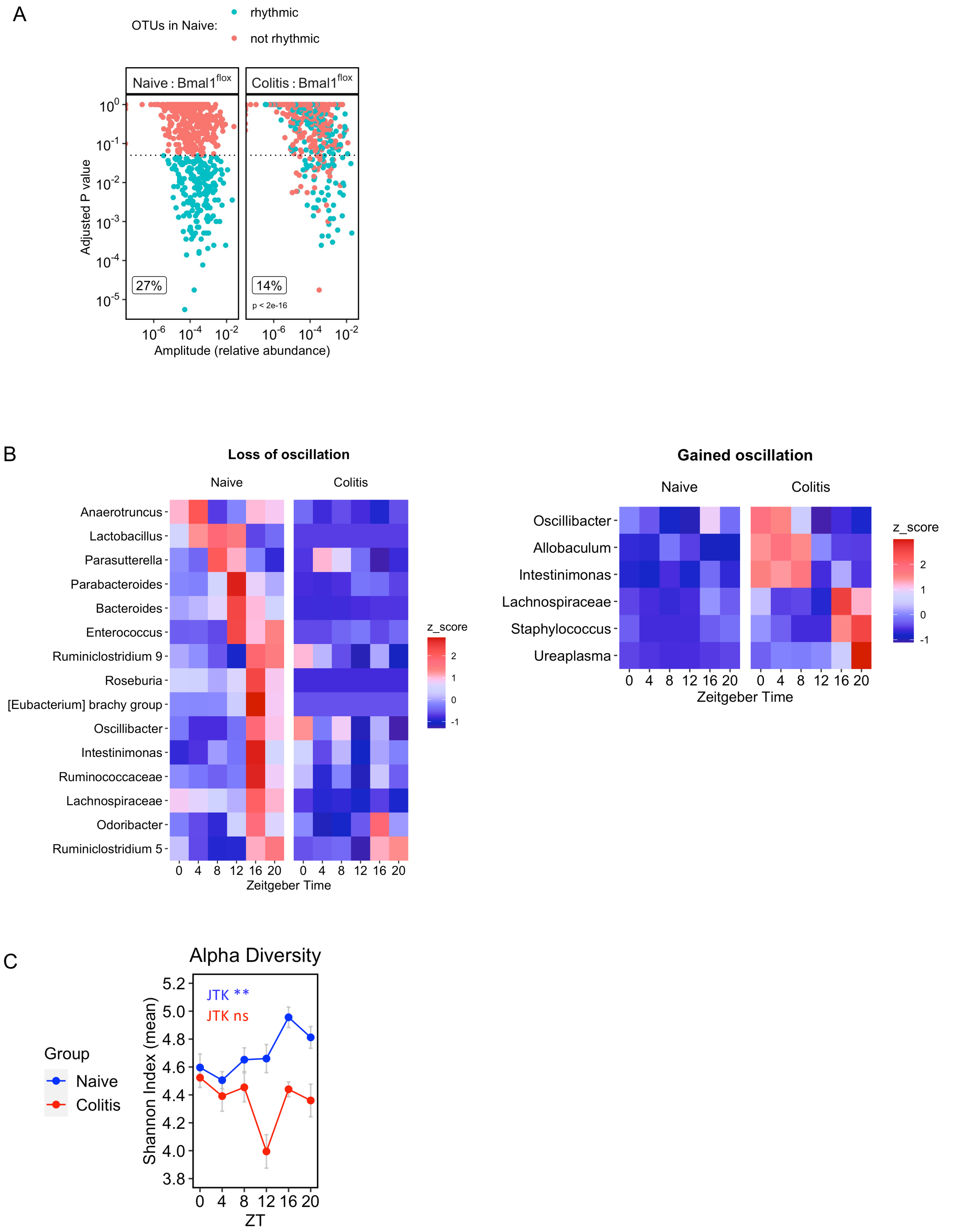

A
OTUs in Naive:
rhythmic
not rhythmic
Naive : Bmal1flox
Colitis : Bmal1flox
Adjusted P value
27%
14%
p < 2e-16
Amplitude (relative abundance)
B
Loss of oscillation
Naive
Colitis
Anaerotruncus
Lactobacillus
Parasutterella
Parabacteroides
Bacteroides
Enterococcus
Ruminiclostridium 9
Roseburia
[Eubacterium] brachy group
Oscillibacter
Intestinimonas
Ruminococcaceae
Lachnospiraceae
Odoribacter
Ruminiclostridium 5
z_score
Zeitgeber Time
Gained oscillation
Naive
Colitis
Oscillibacter
Allobaculum
Intestinimonas
Lachnospiraceae
Staphylococcus
Ureaplasma
z_score
Zeitgeber Time
C
Alpha Diversity
JTK **
JTK ns
Group
Naive
Colitis
Shannon Index (mean)
ZT

**Figure S4: (A)** Rhythmicity in relative abundance of operational taxonomic units (OTUs) in faecal samples harvested around the clock from *Bmal1*$^{flox}$ mice prior to and during (Day 4) DSS colitis (paired samples), n=5/timepoint. **(B)** Heatmaps of comparative rhythmicity for annotated genera. **(C)** Alpha diversity within 16S microbiome sequencing (n=5/treatment/timepoint). Statistics: (A-C) Analysis of rhythmicity by JTK_CYCLE. (A) McNemar test.

### Figure S5

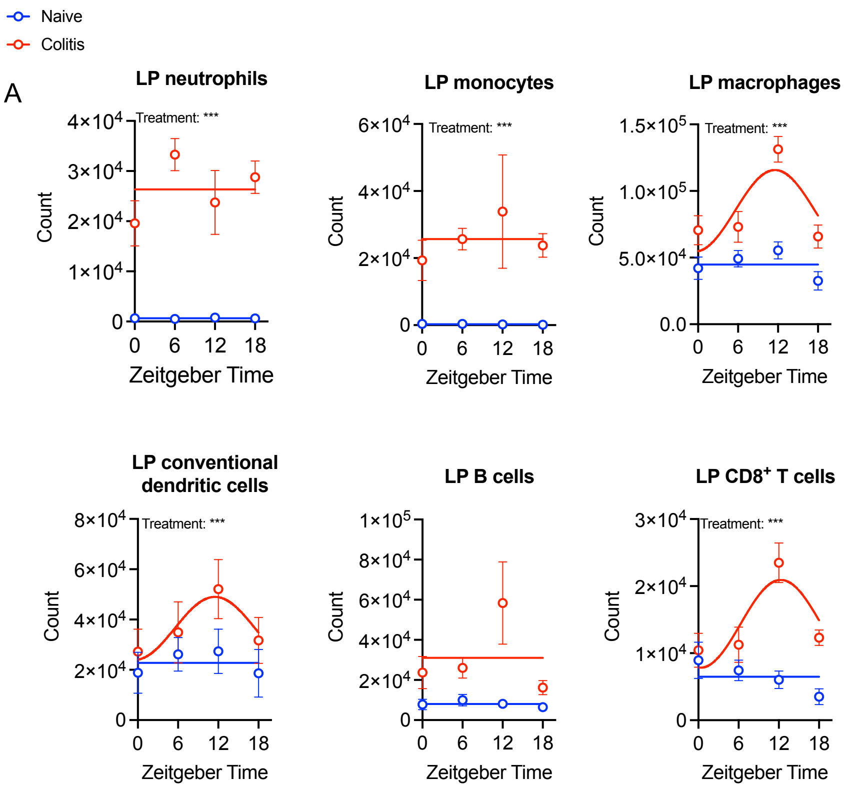
Naive
Colitis
A
LP neutrophils
Treatment: ***
Count
Zeitgeber Time
LP monocytes
Treatment: ***
LP macrophages
Treatment: ***
LP conventional dendritic cells
Treatment: ***
LP B cells
LP CD8+ T cells
Treatment: ***

**Figure S5: (A)** Number of colonic lamina propria (LP) neutrophils (live $CD45^+Ly6G^+$), LP monocytes (live $CD45^+Ly6G^-CD11b^+CD64^+MHCII^-Ly6C^+$), LP macrophages (live $CD45^+Ly6G^-CD11b^+CD64^+MHCII^+Ly6C^-$), LP conventional dendritic cells (live $CD45^+Ly6G^-CD64^-MHCII^+CD11c^+CD103^+CD11b^-$), LP B cells (live $CD45^+$ $CD19^+$) and LP CD8+ T cells (live $CD45^+CD3^+CD8^+$) determined by flow cytometry in wildtype mice (n=5/timepoint/treatment/genotype).

Statistics: (A) Two-way ANOVA with multiple comparisons (Šídák) and nonlinear regression to compare whether best fit is given by horizontal line or sine wave with nonzero baseline, constrained to a wavelength of 24 hours and an amplitude of greater than 0.

### Figure S6

A

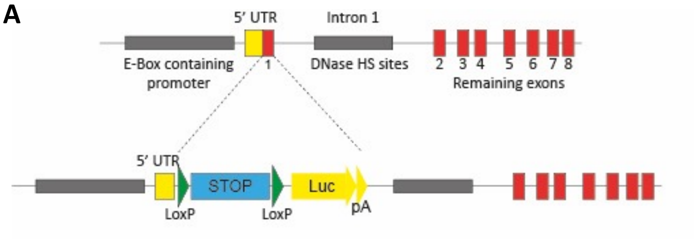
5' UTR
Intron 1
E-Box containing promoter
1
DNase HS sites
2 3 4 5 6 7 8
Remaining exons
5' UTR
STOP
Luc
LoxP
LoxP
pA

B

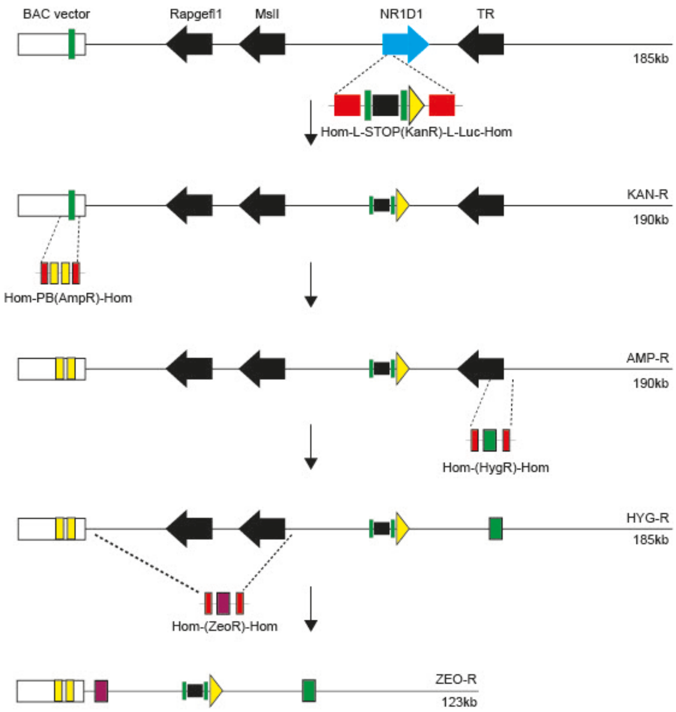
BAC vector
Rapgefl1
Msll
NR1D1
TR
185kb
Hom-L-STOP(KanR)-L-Luc-Hom
KAN-R
190kb
Hom-PB(AmpR)-Hom
AMP-R
190kb
Hom-(HygR)-Hom
HYG-R
185kb
Hom-(ZeoR)-Hom
ZEO-R
123kb

**Figure S6: (A)** Schematic of recombineering cassette to replace NR1D1 exon 1 with a conditional STOP site followed by Luciferase reporter gene. Note the presentation of key gene regulatory features including the upstream promoter, 5'UTR and intron 1 containing putative regulatory elements. **(B)** Overall view of BAC engineering including the integration of the reporter cassette, the PiggyBac ITRs in the BAC vector, and sequential removal of bystander genes.

**Figure S7**

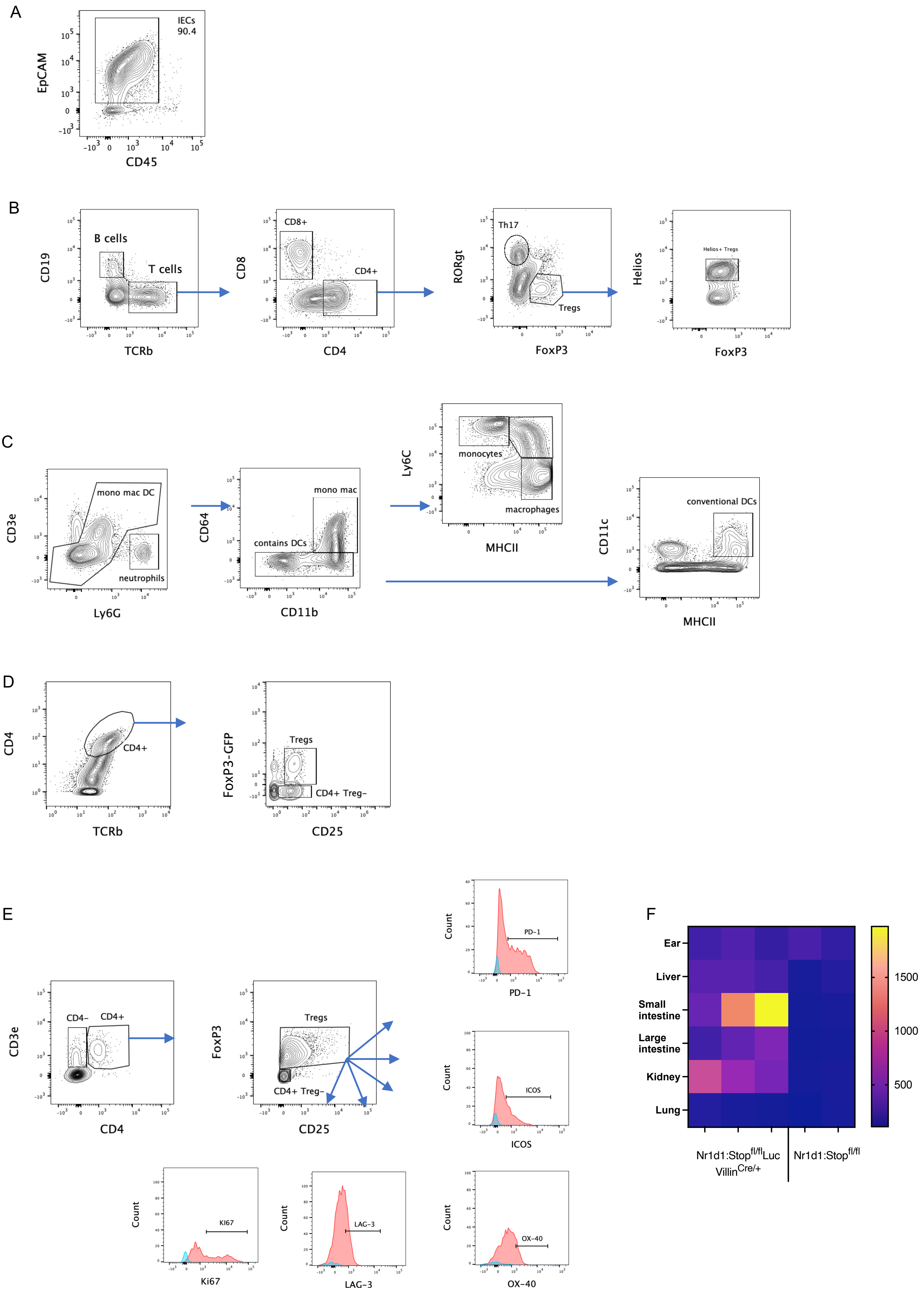
A
IECs
90.4
EpCAM
CD45
B
B cells
T cells
CD19
TCRb
CD8+
CD4+
CD8
CD4
Th17
Tregs
RORgt
FoxP3
Helios+ Tregs
Helios
FoxP3
C
mono mac DC
neutrophils
CD3e
Ly6G
mono mac
contains DCs
CD64
CD11b
monocytes
macrophages
Ly6C
MHCII
conventional DCs
CD11c
MHCII
D
CD4+
CD4
TCRb
Tregs
CD4+ Treg−
FoxP3-GFP
CD25
E
CD4−
CD4+
CD3e
CD4
Tregs
CD4+ Treg−
FoxP3
CD25
PD−1
Count
ICOS
Ki67
LAG−3
OX−40
F
Ear
Liver
Small intestine
Large intestine
Kidney
Lung
Nr1d1:Stop^fl/fl Luc Villin^Cre/+
Nr1d1:Stop^fl/fl
1500
1000
500

**Figure S7: (A)** Flow cytometry data showing purity of the colonic intestinal epithelial cell (IEC) population ($CD45^{-}EpCam^{+}$) as purified by EDTA/DTT incubation. **(B)** Lymphoid gating strategy. Cells pre-gated as single/live/$CD45^{+}$. **(C)** Myeloid gating strategy. Cells pre-gated as single/live/$CD45^{+}$. **(D)** Gating strategy for regulatory T cells ($CD25^{+}GFP^{+}$) sorted from DEREG mice using the BD Influx. Cells pre-gated as single/live **(E)** Gating strategy for regulatory T cell functional panel. Cells pre-gated as single/live/$CD45^{+}$. Blue histogram is fluorescence minus one (FMO), pink histogram is representative sample. **(F)** Heatmap of total bioluminescent flux.
